# BRAKER2: Automatic Eukaryotic Genome Annotation with GeneMark-EP+ and AUGUSTUS Supported by a Protein Database

**DOI:** 10.1101/2020.08.10.245134

**Authors:** Tomáš Brůna, Katharina J. Hoff, Alexandre Lomsadze, Mario Stanke, Mark Borodovsky

**Affiliations:** School of Biological Science, Georgia Tech, Atlanta, GA 30332, USA; Institute of Mathematics and Computer Science, University of Greifswald, 17489 Greifswald, Germany; Center for Functional Genomics of Microbes, University of Greifswald, 17489 Greifswald, Germany; Wallace H Coulter Department of Biomedical Engineering, Georgia Tech and Emory University, Atlanta, GA 30332, USA; School of Computational Science and Engineering, Georgia Tech, Atlanta, GA 30332, USA

## Abstract

Full automation of gene prediction has become an important bioinformatics task since the advent of next generation sequencing. The eukaryotic genome annotation pipeline BRAKER1 had combined self-training GeneMark-ET with AUGUSTUS to generate genes’ coordinates with support of transcriptomic data. Here, we introduce BRAKER2, a pipeline with GeneMark-EP+ and AUGUSTUS externally supported by cross-species protein sequences aligned to the genome. Among the challenges addressed in the development of the new pipeline was generation of reliable hints to the locations of protein-coding exon boundaries from likely homologous but evolutionarily distant proteins. Under equal conditions, the gene prediction accuracy of BRAKER2 was shown to be higher than the one of MAKER2, yet another genome annotation pipeline. Also, in comparison with BRAKER1 supported by a large volume of transcript data, BRAKER2 could produce a better gene prediction accuracy if the evolutionary distances to the reference species in the protein database were rather small. All over, our tests demonstrated that fully automatic BRAKER2 is a fast and accurate method for structural annotation of novel eukaryotic genomes.

## Introduction

Constantly improving next generation sequencing (NGS) technology enables production of a nearly complete eukaryotic genome within weeks or even days. Therefore, accurate automatic methods of genome annotation have been in high demand since the dawn of the NGS era. A self-training algorithm for *ab initio* gene prediction in eukaryotic genomes, GeneMark-ES [1], has accelerated the process of structural annotation for many genome projects [2-7]. Application of NGS to transcript sequencing (RNA-Seq) motivated active development of methods integrating genomic and transcriptomic information. A new automatic algorithm, GeneMark-ET [8], integrated data on spliced aligned RNA-Seq reads into GeneMark-ES.

On a parallel avenue, yet another algorithm, AUGUSTUS [9-14], was demonstrated to be one of the most accurate gene prediction tools [15-17]. AUGUSTUS carried a flexible mechanism for integration of external evidence generated by spliced-aligned RNA-Seq reads or homologous proteins into gene prediction. AUGUSTUS also used this evidence to predict alternative isoforms. Still, for model parameter estimation, AUGUSTUS required an expert curated training set of genes.

It was apparent that a useful automatic tool could be created by combining strong features of GeneMark-ET and AUGUSTUS. A pipeline BRAKER1 was developed and released in 2015 [18] to become a frequently used tool in genome annotation projects [19-24]. BRAKER1 requires availability of RNA-Seq data, however, not all novel genomes are sequenced along with transcriptomes, e.g. within the Earth BioGenome Project [25], or transcriptome coverage may be insufficient.

Here, we introduce BRAKER2 that uses cross-species protein sequence data, readily available for any genome project. Processing of mapped to genome protein sequences has presented a well-known challenge, due to the protein divergence and uneven speed of evolution among protein families. Nonetheless, leveraging of large volumes of proteins, particularly proteins from remotely related species, for improving genome annotation appears to be an important theoretical and practical task. Recently developed GeneMark-EP+ [26] was able to direct self-training by hints created by processing spliced alignments of large numbers of cross-species proteins. It was logical to integrate GeneMark-EP+ and AUGUSTUS in a pipeline relying on already known protein sequences as external evidence.

Several tools were created with a goal to identify a eukaryotic gene structures via spliced alignment of a homologous protein to the genomic locus – PROCRUSTES [27], GenomeThreader [28], ProSplign [29], Scipio [30], Spaln [31], etc. However, it turned out that the accuracy of the spliced alignment deteriorated quickly with increase of the evolutionary distance between two species. GeneMark-EP+ [26] has addressed this problem by the use of spliced alignments of multiple proteins originating from either closely or remotely related species.

The hints to exon borders of a gene are made by the ProtHint pipeline (integral part of GeneMark-EP+) that runs fast homolog detection and protein spliced alignments to the genomic region. ProtHint is using the alignment’s data to score and classify the hints. Homologous proteins, even those from remotely related species, contribute by providing information for hints to the borders of exons encoding evolutionarily conserved protein domains.

Salient features of BRAKER2 are i/ it is fully automatic ii/ it runs massive database search for proteins that are homologous to proteins encoded in the new genome (yet unknown ones) iii/ it processes millions of protein spliced alignments to the genome to generate hints to exon-intron structures, iv/ it integrates sequence composition based and protein alignment based information at all iterative steps of training and gene prediction.

We assessed prediction accuracy of BRAKER2 on well-studied and, arguably, well-annotated genomes of *Arabidopsis thaliana, Caenorhabditis elegans*, and *Drosophila melanogaster*. For tests on additional nine genomes, we selected subsets of annotated genes corroborated by RNA-Seq evidence. We compared performance and accuracy of BRAKER2 to performance and accuracy of MAKER2 [27] with two distinct execution protocols as well as to BRAKER1 [18].

## Materials and Methods

### Materials

For testing BRAKER2, we used genomic sequences and gene annotations of twelve species (Table 1). Among them were the early sequenced model organisms: *A. thaliana, C. elegans*, and *D. melanogaster*. The other nine species were: the plants *Populus trichocarpa, Medicago truncatula, Solanum lycopersicum*, the arthropods *Bombus terrestris, Rhodnius prolixus, Parasteatoda tepidariorum*, and the vertebrates *Tetraodon nigroviridis, Danio rerio* and *Xenopus tropicalis* (Table 1). We used the OrthoDB database [32] as a source of protein data. RNA-Seq data used in runs of BRAKER1 was sampled from the Sequence Read Archive [33] by VARUS [34].

**Table 1:** Genomes used in the tests; asterisks indicate model organisms. An average number of introns per gene was determined with respect to the number of all the annotated genes in the genome. For a gene to be considered complete and canonical, at least one of the gene’s transcripts had to be fully annotated, with the initial coding exon starting with a ‘canonical’ ATG and the terminal coding exon ending with TAA, TAG or TGA.

| Species | Annotation version | Genome size (Mb) | # Genes in annotation | # Introns per gene | % Non-canonical or incomplete genes |
| --- | --- | --- | --- | --- | --- |
| Species with early sequenced genomes |  |  |  |  |  |
| *A. thaliana* * | Tair Araport 11 (Jun 2016) | 119 | 27,445 | 4.9 | 0.3 |
| *C. elegans* * | WormBase WS271 (May 2019) | 100 | 20,172 | 5.7 | 0.2 |
| *D. melanogaster* * | FlyBase R6.18 (Jun 2019) | 138 | 13,929 | 4.3 | 0.3 |
| Other species |  |  |  |  |  |
| Plantae |  |  |  |  |  |
| *P. trichocarpa* * | JGI Ptrichocarpa_533_v4.1 (Nov 2019) | 389 | 34,488 | 4.9 | 0.3 |
| *M. truncatula* * | MtrunA17r5.0-ANR-EGN-r1.6 (Feb 2019) | 430 | 44,464 | 2.9 | 0.0 |
| *S. lycopersicum* | Consortium ITAG4.0 (May 2019) | 773 | 33,562 | 3.5 | 14.5 |
| Arthropoda |  |  |  |  |  |
| *B. terrestris* | NCBI Annotation Release 102 (Apr 2017) | 249 | 10,581 | 7.1 | 4.7 |
| *R. prolixus* | VectorBase RproC3.3 (Oct 2017) | 707 | 15,061 | 4.8 | 34.7 |
| *P. tepidariorum* | NCBI Annotation Release 101 (May 2017) | 1,445 | 18,602 | 7.3 | 18.2 |
| Vertebrata |  |  |  |  |  |
| *T. nigroviridis* | TETRAODON8.99 (Nov 2019) | 359 | 19,589 | 10.4 | 63.8 |
| *D. rerio* * | Ensembl GRCz11.96 (May 2019) | 1,345 | 25,254 | 8.2 | 11.8 |
| *X. tropicalis* * | NCBI Annotation Release 104 (Apr 2019) | 1,449 | 21,821 | 12.1 | 2.4 |

To determine to which degree both predicted and annotated genes covered the sets of universal single copy genes identified by BUSCO protein families, we used the BUSCO database v4 [35].

## Methods

### Description of BRAKER2

First, the *ab initio* gene finder GeneMark-ES [1] completes self-training on a given genome and delivers predicted genes, the initial set of *seed genes*. This step is a part of the internal pipeline ProtHint (described earlier [26]) that executes GeneMark-ES, DIAMOND [36] and Spaln [31] (Fig. 1). The connection between translated seed genes (*seed proteins*) and the genomic loci where the seed genes are residing (*seed regions*) is used in the subsequent steps. The seed proteins make queries for the DIAMOND database search that identifies potentially homologous (target) proteins in a protein database [26]. Next, the newly found target proteins are spliced aligned by Spaln to the genomic *seed regions* where the queries were encoded. The alignments are scored and converted into hints to introns and translation initiation (start codon) and termination sites (stop codon) with scores characterizing the hints’ reliability. A subset of high confidence hints is selected (see Supplementary Materials and [26]). The hints to those exon borders that are also identified by the *ab initio* algorithm (GeneMark-ES) are of special interest. They define *anchored sites* and *regions* for iterative model training of GeneMark-EP [26]. Moreover, the high confidence hints to exon borders are enforced in GeneMark-EP+ predictions [26].

**Figure 1:**
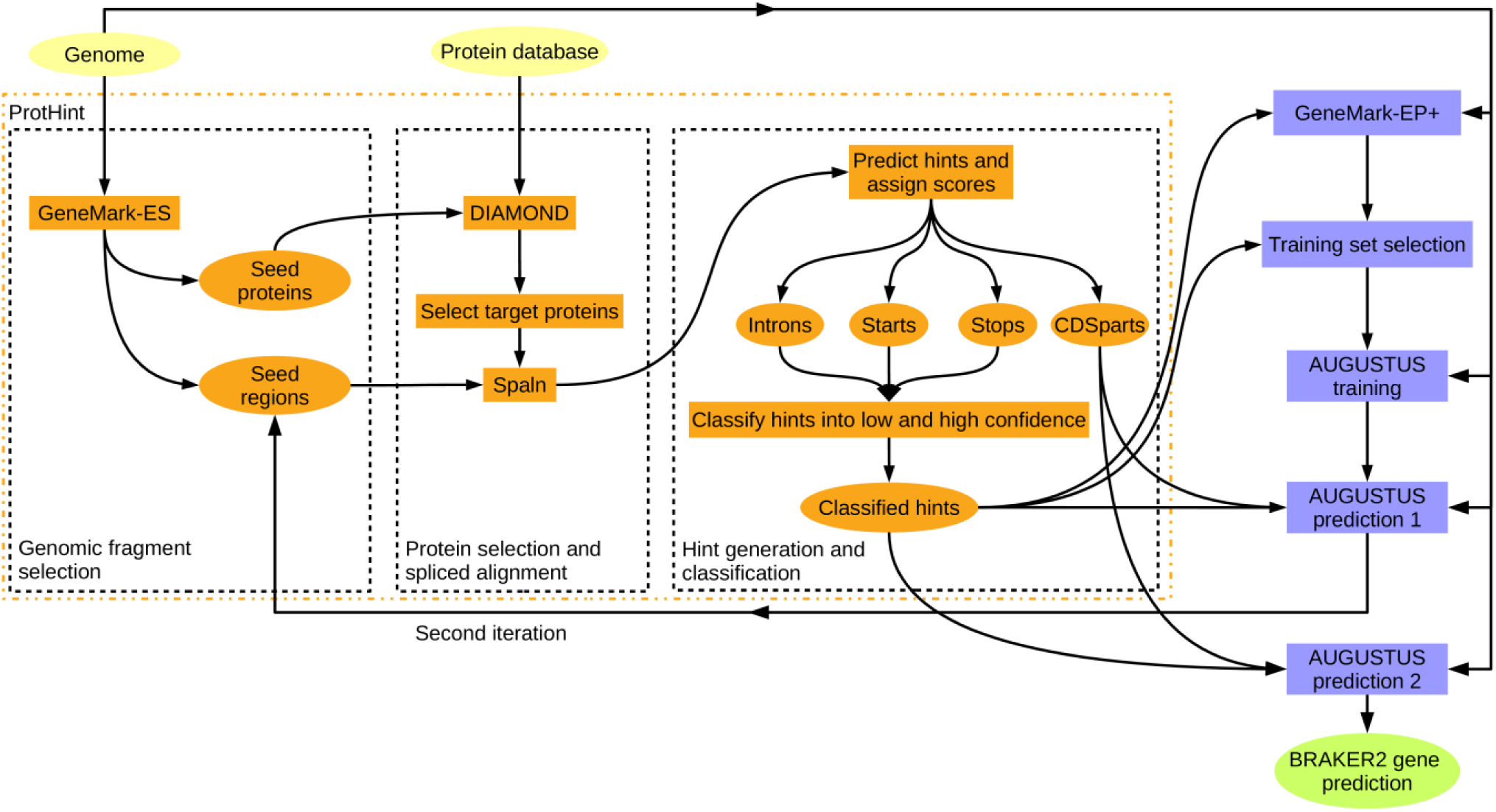
Flowchart of the BRAKER2 pipeline. Input files are shown as yellow ovals, the final output file is shown as green oval. Tools, processes and content types within the ProtHint pipeline are shown in orange, while other components of BRAKER2 are shown in blue.

In BRAKER2, in addition to the hint scheme implemented in GeneMark-EP+, ProtHint makes hints called *CDSpart chains*. This type of hints helps to combine exons predicted by AUGUSTUS into a single transcript. The *CDSpart chain* is defined by a spliced alignment of the highest scoring target protein to the seed region.

From the whole complement of genes predicted by GeneMark-EP+, we select a set of *anchored genes* – the single-exon genes with hints matching start- and stop-codons as well as the multi-exon genes with all introns supported by hints. These genes make a training set for AUGUSTUS that, in turn, enforces *the high confidence* hints in gene predictions. The *CDSpart chain* hints and non-high-confidence hints are processed for integration into AUGUSTUS gene prediction as well (Supplementary Materials, Sections 1.2 and 1.3). Note that in the regions lacking extrinsic evidence, GeneMark-EP+ and AUGUSTUS predict genes in an *ab initio* mode, only.

BRAKER2 runs in two major iterations (Fig. 1). The first one starts with the seed genes predicted by GeneMark-ES [1]. Seeds for some true genes might be missed at this stage; however, they could be recovered in the second BRAKER2 iteration that uses the genes predicted in the first iteration as seed genes. In the second iteration, ProtHint runs the database search only for the newly added seed genes and merges the newly defined hints with the hints from the first iteration. Then, AUGUSTUS uses the models trained in the first iteration along with the updated protein hints to predict the final set of genes. The second BRAKER2 iteration has fewer steps and runs faster than the first iteration.

### Accuracy assessment

#### Selection of protein data sets and test sets of annotated genes

A test of BRAKER2 on a well-studied genome should utilize a set of cross-species proteins that imitates a protein set available for running BRAKER2 on a newly sequenced genome. Proteins that originate from the most evolutionarily close species are expected to be most informative for the BRAKER2 algorithm. Therefore, a meaningful characteristic of a selected set of reference proteins is the least evolutionary distance from the reference genomes to the genome in the test.

To make these selections for *A. thaliana, C. elegans*, and *D. melanogaster*, we have started from large clades (Plantae, Metazoa, Arthropoda, respectively) and have created three sets of proteins for each species by excluding either (i) proteins from the given species per se, (ii) proteins from all species of the same *family*, (iii) proteins from all species of the same *order*. For the other nine species, we also have defined large clades and then have only used partitions of type (iii) (Table 2).

**Table 2:** Numbers of species in the clades of OrthoDB v10. Bold font numbers mark the largest OrthoDB clade used to support gene predictions for a given species. The numbers shown in blue give the numbers of species in the smaller clades removed from the largest OrthoDB segment in the tests described below. Species whose proteins are not present in the current OrthoDB version are marked with asterisks.

| Species | # of species in the OrthoDB clade |  |  |  |  | Name of the largest OrthoDB segment | # of proteins in the OrthoDB segment |
| --- | --- | --- | --- | --- | --- | --- | --- |
|  | Family | Order | Class | Phylum | Kingdom |  |  |
| *A. thaliana* | **8** | **10** | - | 100 | **117** | Plantae | 3,510,742 |
| *C. elegans* | **3** | **5** | 6 | 7 | **448** | Metazoa | 8,266,016 |
| *D. melanogaster* | **20** | **56** | 148 | **170** | - | Arthropoda | 2,601,995 |
| *P. trichocarpa* * | 5 | **5** | - | 100 | **117** | Plantae | 3,510,742 |
| *M. truncatula* | 10 | **10** | - | 100 | **117** | Plantae | 3,510,742 |
| *S. lycopersicum* | 10 | **11** | - | 100 | **117** | Plantae | 3,510,742 |
| *B. terrestris* * | 7 | **40** | 148 | **170** | - | Arthropoda | 2,601,995 |
| *R. prolixus* | 1 | **16** | 148 | **170** | - | Arthropoda | 2,601,995 |
| *P. tepidariorum* | 1 | **2** | 10 | **170** | - | Arthropoda | 2,601,995 |
| *T. nigroviridis* * | 1 | **1** | 50 | **246** | - | Chordata | 5,003,104 |
| *D. rerio* | 5 | **5** | 50 | **246** | - | Chordata | 5,003,104 |
| *X. tropicalis* | 2 | **3** | 3 | **246** | - | Chordata | 5,003,104 |

In the tests done with 12 eukaryotic genomes (Table 1), we used as ‘gold standards’ either whole genome annotation (*A. thaliana, C. elegans*, and *D. melanogaster*) or annotation of sets of complete multi-exon genes with all introns fully supported by mapped RNA-seq reads.

Protein-coding gene prediction accuracy was defined at exon and gene levels by values of sensitivity (Sn), specificity (Sp) as well as their harmonic mean (F1), with the definitions given in the Supplementary Materials. At the gene level, in presence of alternative splicing, a gene was considered to be predicted correctly if the predicted CDS matches precisely a CDS of one of the annotated transcript isoforms.

#### Use of universal single copy genes from BUSCO families

The BUSCO metrics is supposed to evaluate the completeness of a genome assembly and annotation; it is based on collections of single copy genes expected to be present in a particular lineage [35]. The ‘BUSCO genes’ may constitute less than 5% of genes in the genome, nonetheless, this approach is practical for novel genomes given its relatively easy application. We used the BUSCO metrics to characterize gene or protein sets predicted by BRAKER2 in several genomes.

While, the BUSCO metrics give an idea on the gene prediction algorithm’s Sn value, it does not quantify the algorithm’s tendency to predict false positives (the Sp value). Moreover, since the BUSCO based method relies on HMMER3 [37] search for detecting homologs of the BUSCO proteins, it does not discriminate between precisely and approximately predicted exon-intron structures. Therefore, the BUSCO metrics are less precise in assessment of accuracy of gene prediction than the methods comparing coordinates of predicted and annotated genes and computing Sn, Sp and F1 values.

#### Testing MAKER2

The MAKER2 genome annotation pipeline can combine information from several sources, such as *ab initio* gene predictions, mapped RNA-Seq reads as well as alignments of proteins to the genome [38-40].

For our tests, we have chosen genomes of *A. thaliana, C. elegans*, and *D. melanogaster*, arguably the best annotated genomes among the genomes that we have considered. Also, for each species, we have used the relevant segment of the OrthoDB database described above, with exclusion of species of the same taxonomic *order*.

All the components of the MAKER2 pipeline, e.g. repeat annotation or training of gene finders, have been executed in *de novo* mode, i.e. each of the three genomes was considered to be a “novel” one.

By design, the protein mapping in MAKER2 is much slower than protein mapping done by ProtHint in BRAKER2, therefore, we have further limited each of the three OrthoDB partitions to randomly selected ten species (Table S3). We have used two MAKER2 execution protocols (described in detail in Supplementary Materials). In the first protocol recommended by the authors [40], the protein spliced alignments have been used to create training sets for AUGUSTUS and SNAP. The final gene predictions have been made by combining predictions of self-training GeneMark-ES with ones from AUGUSTUS and SNAP both using the protein derived hints (Fig. S5A). We have introduced the second training protocol, somewhat similar to the one of BRAKER2, in which protein spliced alignments and GeneMark-ES predictions have been used to create a training set for AUGUSTUS. The final gene predictions have been made by only two gene finders, GeneMark-ES, and AUGUSTUS with hints (Fig. S5B).

MAKER2 offers two modes of gene prediction: to only get predictions supported by external evidence or to add predictions generated without support. Given that the set of proteins has provided support for a limited number of genes (Table S7), we have executed MAKER2 in the second mode, the one producing higher Sn values.

The repeat masking for both BRAKER2 and MAKER2 has been done with the same genome specific repeat library (generated by RepeatModeler). Training and predictions have been done on a repeat-masked sequence. However, BRAKER2 and MAKER2 have different methods for processing repeat-masked sequences (see Supplementary Materials).

#### Testing BRAKER1

BRAKER1 is a genome annotation pipeline that combines self-training GeneMark-ET with AUGUSTUS. External evidence in a form of short RNA-Seq reads to genome alignments is used to generate hints to intron borders [12]. BRAKER1 and BRAKER2 use conceptually similar features, such as anchored elements of exon-intron structure.

BRAKER1 has been run on the genomes of *A. thaliana, C. elegans*, and *D. melanogaster* with hints originating from RNA-Seq reads sampled by VARUS [34] from the NCBI Sequence Read Archive [33]. VARUS used HISAT2 [41] for mapping RNA-Seq reads to genomic sequences (Supplementary Materials, section 1.10).

## RESULTS

### Assessment of BRAKER2 accuracy on genomes of *A. thaliana, C. elegans*, and *D. melanogaster* and comparison with BRAKER1

The accuracy of BRAKER2 was determined at exon and gene level (Figs. 3-4). The exon level Sn and Sp showed the following patterns (Fig. 3). In comparison with accuracy reached by the *ab initio* GeneMark-ES, BRAKER1 clearly improved both Sn and Sp values. On the other hand, BRAKER2 delivered better results than BRAKER1 when the set of reference proteins was the largest for each genome (excluding only proteins from the same species). For the two smaller protein sets for each species: excluding the same family or the same order, the results of comparison were mixed. BRAKER2 was better both times for *A. thaliana*, but not for *D. melanogaster*, and especially not for *C. elegans* (Fig.3).

**Figure 2:**
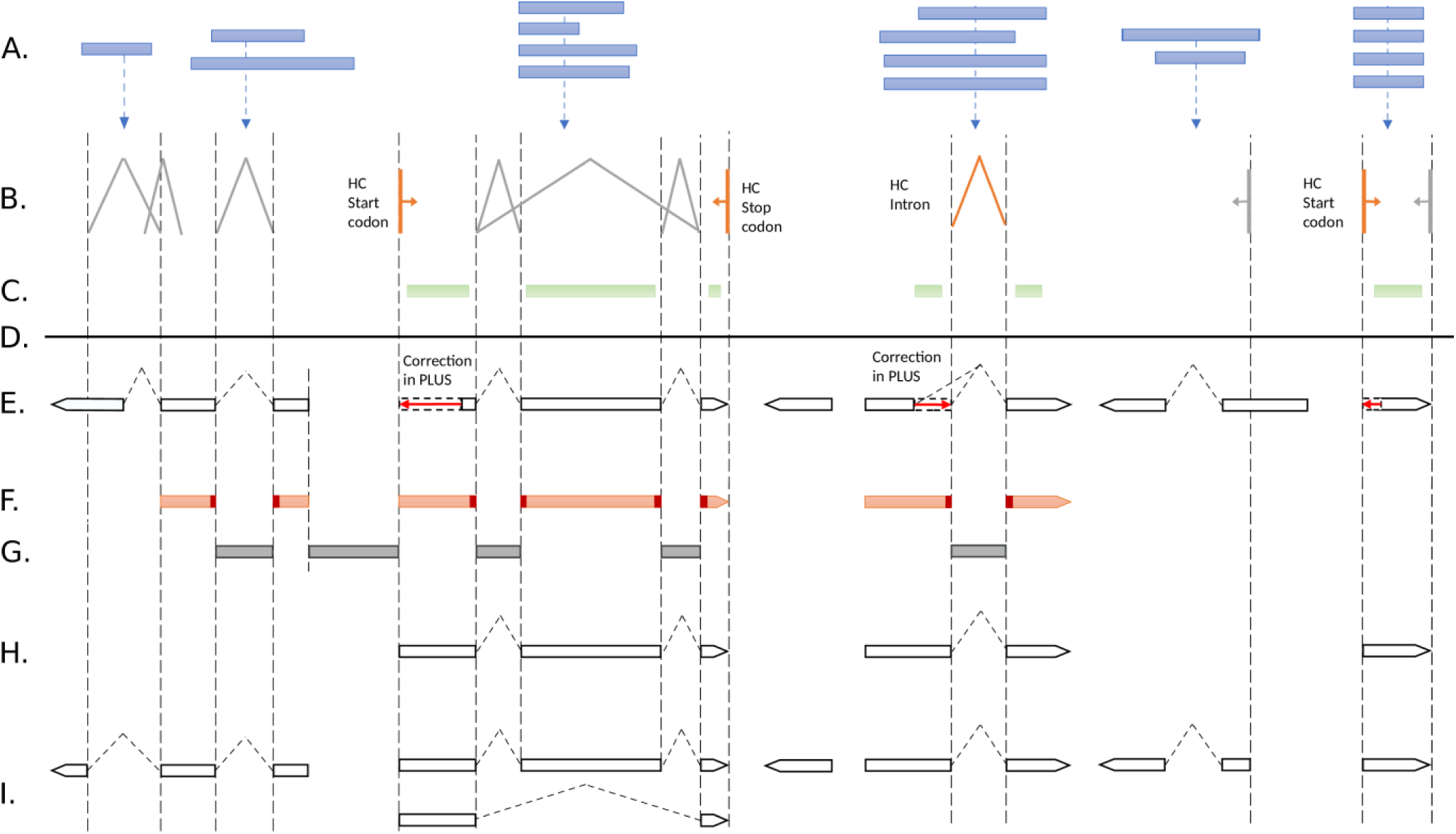
Evidence integration in BRAKER2. A. Target proteins; B. Introns, gene start and stop sites defined by spliced alignments of target proteins to genome; C. CDSpart chains; D. Genome sequence; E. Genes predicted by GeneMark-EP+ at a given iteration. The high confidence hints are enforced (red arrows); F. Anchored sites, the splice sites and gene ends predicted *ab initio* and corroborated by protein hints; G. Anchored introns and intergenic sequences bounded by anchored gene ends are selected into training of non-coding sequence model for GeneMark-EP+; H. Anchored multi-exon and single exon genes predicted by GeneMark-EP+ and selected for training AUGUSTUS; I. Transcripts predicted by AUGUSTUS with support of an external evidence.

**Figure 3:**
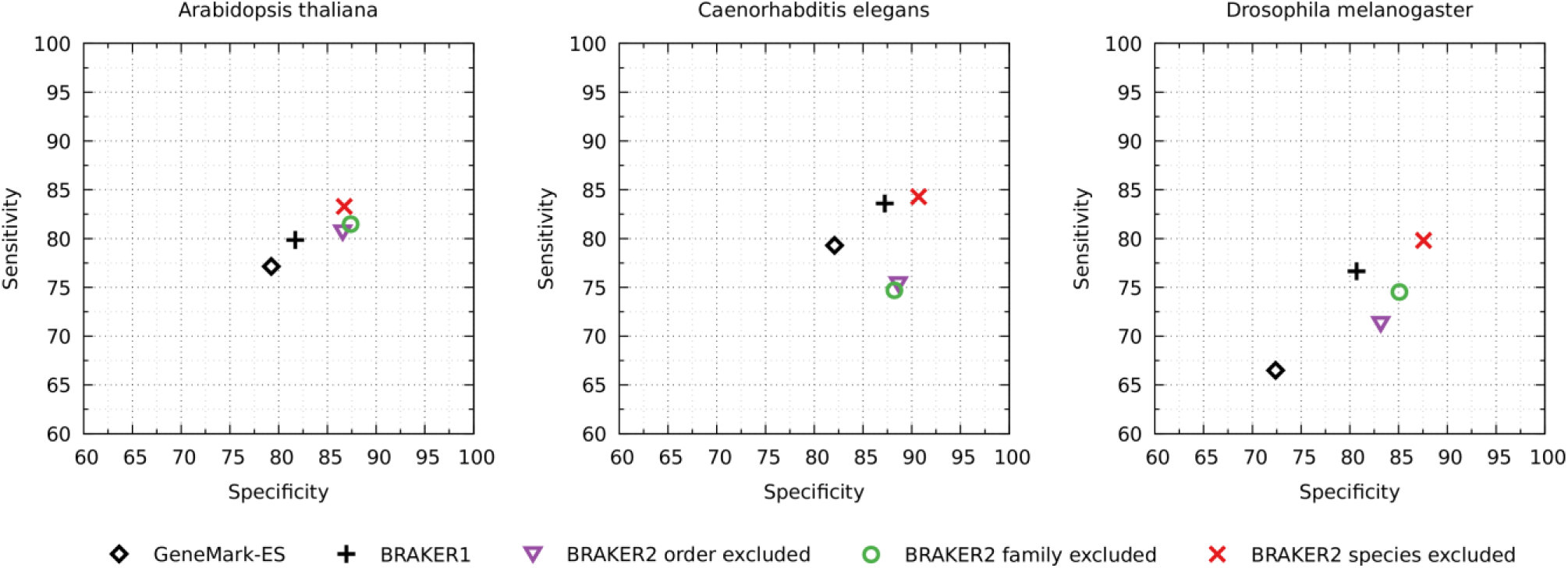
Exon level Sn and Sp are shown for each species for the three test runs of BRAKER2, the runs of GeneMark-ES and BRAKER1 with RNA-Seq support. BRAKER2 for each species was run with a relevant set of OrthoDB that did not include proteins i/ of the same species, ii/ of all species of the same family, iii/ of all species of the same order.

The pattern of accuracy change at exon level was mainly translated into the accuracy at gene level (Fig. 4, Tables S4, S5, S6). The order of the tested annotation tools from the lower to higher accuracy became less ambiguous, since the vectors (Sp, Sn) were lined up along a diagonal. One of the differences in comparison with the patterns at exon level was that BRAKER2 outperformed BRAKER1 on the genome of *D. melanogaster* when the reference proteins were comprised from the phylum *Arthropoda* excluding the species of the same family. The F1 values corresponding to Fig. 4 are listed in Tables S4, S5, S6.

**Figure 4:**
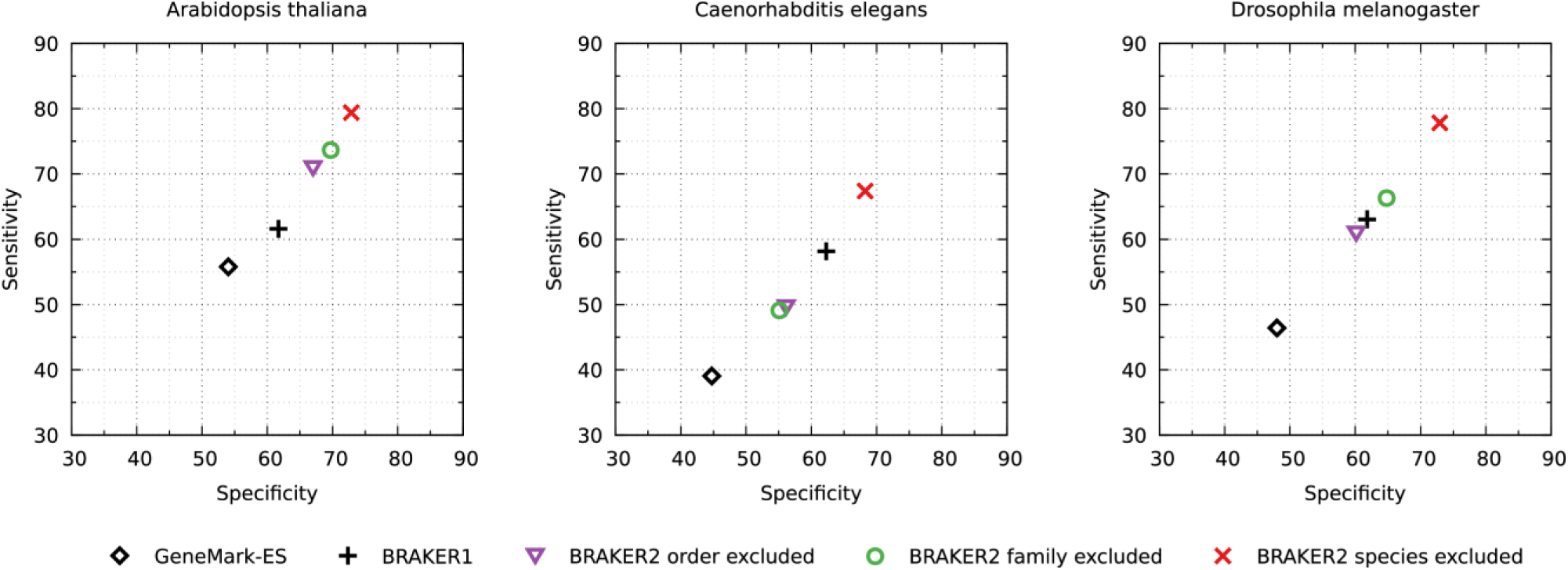
Gene level Sn and Sp observed in the same tests as described in the legend for Fig. 3.

### Assessment of the BRAKER2 accuracy on the nine more genomes

Given that *A. thaliana, C. elegans*, and *D. melanogaster* were subjects of the pilot genome sequencing projects, we used longtime curated annotations of these three genomes as large test sets.

Our approach to conducting tests on genomes of the other nine species (Table 3, the species names shown in blue color) was a bit different. We did not have confidence that comparison with the whole genome annotation would give a fair assessment of the method accuracy. To clarify this issue, we compared BRAKER2 gene predictions for the *R. prolixus* genome with the current genome annotation (Table 1). The Sn value at gene level was 13.2% (Table 3). However, we saw an increase of the Sn value to 44.5% when the comparison used a set of complete multi-exon genes with all introns supported by at least one mapped RNA-Seq read (a 26.4% subset of all multi-exon genes). A relatively large difference between gene level Sn measured on the whole genome and the one computed on ‘curated’ gene sets was characteristic for these nine genomes except for *P. trichocarpa* and *X. tropicalis*.

**Table 3:** BRAKER2 gene prediction sensitivity (Sn) at gene and exon level. The test sets were made from (a) all annotated multi-exon genes and (b) reliable genes: annotated complete multi-exon genes with all introns supported by RNA-Seq reads sampled by VARUS [30].

| Species | Gene Sn |  | Exon Sn |  | % Reliable Genes |
| --- | --- | --- | --- | --- | --- |
|  | All | Reliable | All | Reliable |  |
| *A. thaliana* | 70.2 | 78.8 | 81.5 | 87.9 | 83.5 |
| *C. elegans* | 49.8 | 57.8 | 75.7 | 81.0 | 81.1 |
| *D. melanogaster* | 59.5 | 61.6 | 71.9 | 74.4 | 93.2 |
| *P. trichocarpa* | 69.3 | 76.4 | 86.2 | 90.4 | 84.6 |
| *M. truncatula* | 48.3 | 63.2 | 82.7 | 90.0 | 69.6 |
| *S. lycopersicum* | 40.7 | 68.0 | 78.5 | 92.1 | 54.4 |
| *B. terrestris* | 45.7 | 56.7 | 74.6 | 79.5 | 75.1 |
| *R. prolixus* | 13.2 | 45.5 | 61.4 | 80.2 | 26.4 |
| *P. tepidarium* | 24.6 | 40.2 | 67.9 | 79.9 | 50.6 |
| *T. nigroviridis* | 10.4 | 67.7 | 60.6 | 89.5 | 11.2 |
| *D. rerio* | 39.1 | 50.3 | 75.6 | 86.3 | 70.8 |
| *X. tropicalis* | 38.9 | 46.3 | 75.3 | 80.0 | 74.8 |

As expected, the Sn values for *A. thaliana, C. elegans*, and *D. melanogaster*, determined for the whole complements of genes and for RNA-Seq supported complete multi-exon genes, differed by smaller margins in comparison with the other nine genomes (Table 3).

Therefore, we assume that the test sets supported by RNA-Seq provide more objective values for the algorithm accuracy. Particularly, we observed that for the three arthropods the exon Sn values were near 80%, for the three vertebrates they were in the range from 80 to 87%, and for the three plants they were around 90%, the highest (Table 3).

Among the genes predicted by BRAKER2 in the nine genomes, we identified genes encoding proteins that belonged to the BUSCO protein families. Next, we determined the percentages of such genes with respect to the complete species-specific set of the BUSCO protein families expected to reside in a particular genome. The same computation was done for the genes present in the reference genome annotation (Fig. S2).

In the plant and arthropods genomes, BRAKER2 missed ∼3% or less of the BUSCO families. Also, fewer BUSCO families were missed among BRAKER2 genes than in the current annotation of genomes of *M. truncatula, S. lycopersicum, P. tepidariorum* and *R. prolixus*

In the vertebrate genomes the percentage of BUSCO families missed by BRAKER2 was: ∼12% in *T. nigroviridis*, 5% in *D. rerio*, 9% in *X. tropicalis* while the annotations missed ∼12%, 3% and 3%, respectively, of representatives of the BUSCO families.

### Dynamics of the accuracy change in internal steps of BRAKER2

We observed a steady increase from one to another step of the BRAKER2 pipeline for *D. melanogaster, A. thaliana* and *C. elegans* (Table S12). For instance, at gene level the F1 value for *D. melanogaster* increased from GeneMark-ES to GeneMark-EP+ by 17.1 percentage points. Runs of AUGUSTUS with hints added 8.2 percentage points in the first iteration, and 1.1 percentage points in the second iteration.

For the F1 value at the exon level, the numbers of increase were 8.8, 4.6 and 0.4 percentage points, respectively.

### Selection of training genes

In BRAKER2, we optimized selection of genes for training AUGUSTUS. Instead of a complete set of genes predicted by GeneMark-EP+, we used only fully anchored genes. In *A. thaliana, C. elegans*, and *D. melanogaster* the use of these narrower sets improved the gene level F1 values of the *ab initio* gene prediction by AUGUSTUS by two to five percentage points (Table S8). The *ab initio* gene prediction accuracy of AUGUSTUS was chosen as a criterion here instead of accuracy of the full BRAKER2 because accurate hints could overshadow the effects of training.

The use of anchored genes for the AUGUSTUS training had a stronger effect for large genomes where the difference in F1 value at exon level for *D. rerio* reached ∼10 percentage points (Table S8).

### Repeat masking

Repetitive sequences (interspersed repeats and low complexity sequences) were identified by RepeatModeler [42] and RepeatMasker [43]. A run of RepeatModeler on a whole genome produced a repeat library. Next, the locations of repeats were identified and soft-masked by RepeatMasker.

Repeat masking by RepeatModeler/RepeatMasker with default settings was sufficient to achieve high prediction accuracy in all the tested genomes except for *X. tropicalis*. Its genome contained a large number of long tandem repeats (∼60Mb in total) identified by a run of Tandem Repeats Finder (TRF) with *maximum repeat period size* = 500 [44]. The presence of the tandem repeats with elevated GC content (Figure S4), when left unmasked, caused GeneMark-ES, running in the first step of BRAKER2, to converge to an incorrect model. This model made GeneMark-ES predict a majority of coding exons (93%) in the GC-rich regions of long tandem repeats and to poorly predict the true genes.

When we applied the non-standard mode of masking by TRF to other genomes, no significant change in the BRAKER2 prediction accuracy was observed (data not shown). This was not surprising because the additional repeats found by TRF in genomes other than *X. tropicalis* were short in size and similar in GC content to the rest of each genome.

### Assessment of accuracy of MAKER2 and comparison with BRAKER2

The sets of genes predicted by MAKER2 were compared to the reference annotations of the three genomes, *A. thalia*na, *C. elegans*, and *D. melanogaster* (Table 4).

**Table 4:**
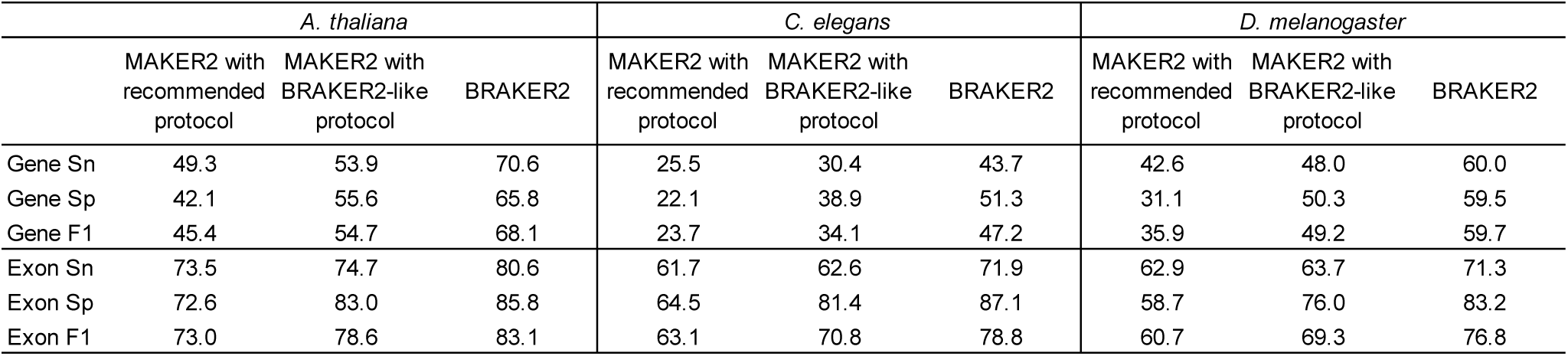
Prediction accuracy of MAKER2 and BRAKER2

|  | *A. thaliana* |  |  | *C. elegans* |  |  | *D. melanogaster* |  |  |
| --- | --- | --- | --- | --- | --- | --- | --- | --- | --- |
|  | MAKER2 with recommended protocol | MAKER2 with BRAKER2-like protocol | BRAKER2 | MAKER2 with recommended protocol | MAKER2 with BRAKER2-like protocol | BRAKER2 | MAKER2 with recommended protocol | MAKER2 with BRAKER2-like protocol | BRAKER2 |
| Gene Sn | 49.3 | 53.9 | 70.6 | 25.5 | 30.4 | 43.7 | 42.6 | 48.0 | 60.0 |
| Gene Sp | 42.1 | 55.6 | 65.8 | 22.1 | 38.9 | 51.3 | 31.1 | 50.3 | 59.5 |
| Gene F1 | 45.4 | 54.7 | 68.1 | 23.7 | 34.1 | 47.2 | 35.9 | 49.2 | 59.7 |
| Exon Sn | 73.5 | 74.7 | 80.6 | 61.7 | 62.6 | 71.9 | 62.9 | 63.7 | 71.3 |
| Exon Sp | 72.6 | 83.0 | 85.8 | 64.5 | 81.4 | 87.1 | 58.7 | 76.0 | 83.2 |
| Exon F1 | 73.0 | 78.6 | 83.1 | 63.1 | 70.8 | 78.8 | 60.7 | 69.3 | 76.8 |

When we used the recommended MAKER2 protocol (Fig. S5a), the accuracy was significantly lower than the one of BRAKER2 ran with support of the same reference proteins. Particularly, the exon F1 measure was lower for *A. thaliana, C. elegans* and *D. melanogaster* by 10.1, 15.7 and 16.1 percentage points, respectively (Table 4). The F1 gap between MAKER2 and BRAKER2 was reduced by running the second protocol (Fig. S5b) to 4.5, 8.0 and 7.5 percentage points, respectively (Table 4). The large improvement in the observed Sp value could be explained by the exclusion of SNAP that generated an elevated number of false positive predictions (Tables S9, S10).

The runtimes of BRAKER2 and MAKER2 in our experiments were difficult to compare directly. We executed MAKER2 in the MPI mode on a computational cluster with 96 CPUs. The runtime of MAKER2 (∼10h) using proteins from 10 species was comparable to a time needed for a run of BRAKER2 with proteins from 443 species executed on a single node with 8 CPUs.

## DISCUSSION

### Genome size

We evaluated accuracy of BRAKER2 on genomes that varied in size from 100 Mb (*C. elegans*) to 1.4 Gb (*X. tropicalis*). Interestingly, the observed exon level Sn value remained at about the same level of 80-90% for both shorter and longer genomes, when computed on test sets of ‘reliable genes’ (Table 3). However, the gene level Sn values showed noticeable negative correlation with genome size as determined on the same test sets.

All the genomes used in this study had relatively homogeneous nucleotide compositions. Current versions of the algorithms used in BRAKER2 employed a single set of species-specific models. Accuracy of BRAKER2 would drop down on genomes with heterogeneous composition, such has human (mammalian) or rice (grasses) where several models reflecting heterogeneous genome composition are necessary.

### Size of the protein database and distribution of evolutionary distances to the reference proteins

The accuracy of a gene finding algorithm, which utilizes cross-species proteins mapping, depends strongly on the evolutionary distance between the species [10, 26, 45]. This distance can play a more significant role than the overall number of proteins. For example, the largest set of proteins was available for *C. elegans* (Table 2), still the accuracy of BRAKER2 at the gene level was lower than for *A. thaliana* and *D. melanogaster* (Fig. 4).

The main pattern observed for each species at the gene level was that the increase of evolutionary distance to the closest relative led to a decrease of the BRAKER2 accuracy. The highest accuracy was observed when the species of the same family were present (species excluded), next down was when the species of the order were present (family excluded) and the next when the species of the same taxonomic class were present (order excluded).

Notably, for all three species, when the largest number of reference proteins was available, only exempting proteins that originated from the tested genome, BRAKER2 was more accurate than BRAKER1 with the largest possible RNA-Seq support (Fig. 4).

Of course, there could be a situation that a species with sequenced genome exists at a very close distance, e.g. when the two genomes’ average nucleotide identity (ANI) is at the level of ∼99%. In this case, methods that merely focus on annotation transfer from one genome to another could be more efficient than BRAKER2 assuming that the reference annotation is of high quality. Otherwise, use of BRAKER2 will be a reasonable choice since BRAKER2 is robust to a presence of errors in proteins of the species taken as a reference.

### General issues with compiling training gene sets

*Ab initio* self-training algorithms were shown to deliver high annotation accuracy, but, arguably, training on a sufficiently large set of manually selected gene structures (a supervised training) could outperform, albeit slightly, a self-training algorithm. For instance, AUGUSTUS trained on a randomly selected set of genes annotated in a well-studied genome slightly outperforms AUGUSTUS trained by BRAKER2 in *ab initio* mode. It is important to emphasize, that such an idealistic condition (random sampling from a 100% correct annotation) would be almost never encountered in practice.

The attempts to create a large enough training set were made by approaches centered around mapping of highly conserved cross-species proteins. Nonetheless, the difficulty of getting unbiased parameter sets presented a challenge that was not met until now [46].

As yet another attempt, in the course of this project, we have used the BUSCO protein families to generate training sets of genes for *A. thaliana, C. elegans*, and *D. melanogaster*. Still, with parameters trained on the ‘BUSCO genes’, gene prediction accuracy was lower than the accuracy achieved by BRAKER2 (Table S11).

The new approach used in BRAKER2 has led to a significant increase in the size of the gene set supported by protein evidence. For almost all the species selected for our tests, more than four thousand gene structures were selected into training sets.

### Improvement in the method for generation of external evidence

BRAKER2 is using a new procedure for creating protein hints. The protein mapping pipeline, ProtHint, makes sets of hints with higher and lower confidence. All hints contribute to generation of anchored genes used in training. GeneMark-EP+ is enforcing high confidence hints in the prediction step. In turn, AUGUSTUS utilizes low and high confidence hints at the prediction step along with information about the hints’ connections within a putative transcript (CDSpart chain). The flexible use of hints leads to an increase in accuracy of BRAKER2 (Table S13). Particularly, BRAKER2 appears to be a useful tool for annotation of genomes of deep branching species, since BRAKER2 is tuned up to generate accurate hints by use of proteins from remotely related species.

### BRAKER2 iterations

The number of hints generated by the ProtHint pipeline depends on the number of genes predicted by GeneMark-ES. A solid performance of GeneMark-ES was demonstrated [8, 26], but any *ab initio* gene finder may miss genes. Missed genes would translate in BRAKER2 into missed protein hints to the corresponding genomic loci. The second iteration of BRAKER2 finds hundreds of missed genes and leads to an increase of gene prediction accuracy (Table S12). Still, the effect of the second iteration on the overall accuracy of BRAKER2 is relatively small. Another though computationally expensive way to recover some missed genes is to align proteins from a protein database to the 6-frame translated genomic DNA [39]. Our experiments with such an approach (data not shown) did not lead to a better Sn value than the iterative procedure of BRAKER2.

### Completeness of BRAKER2 gene predictions

In assessments of completeness of the BRAKER2 predicted gene sets from BUSCO families we used a standard method offered by the BUSCO tools [35]. With exception of *X. tropicalis*, the predicted sets of genes were comparable to, or, even more complete than the sets of ‘BUSCO genes’ present in the reference genome annotations. Notably, some BUSCO families were built from species within the same taxonomic order (e.g. *Hemiptera* order of *R. prolixus* or *Solanales* order for *S. lycopersicum*). At the same time, the only protein input to BRAKER2 were proteins of the species *outside* of the corresponding taxonomic order.

A lower level of accuracy of BRAKER2 for *X. tropicalis* could be explained by the insufficient number of homologous proteins. After the exclusion of proteins from the *Anura* taxonomic order from the OrthoDB partition, no proteins from the *Amphibia* taxonomic *class* were left among input proteins (Table 2). Also, a general cause for missing the ‘BUSCO genes’ could be inaccuracy of the *de novo* repeat masking. For example, we observed that among annotated genes missed by BRAKER2 in the *P. trichocarpa* genome, more than half were genes partially masked due to overlaps with long repeats (> 1,000 nt).

### Comparison with MAKER2

The difference in performance and accuracy of MAKER2 and BRAKER2 observed in our experiments was quite large despite the attempt to find a way to improve the MAKER2 protocol (Table 4). The difference in outcomes could be caused by the differences in training, processing repeats, ways of generating and selecting external evidence as well as combining predictions into the final annotation. Many of these differences were presented in sufficient details in descriptions of the protocols used for running MAKER2 (Supplementary Materials).

While MAKER2 uses the *ab initio* self-training algorithm GeneMark-ES, BRAKER2 uses the recent self-training GeneMark-EP+ algorithm that integrates protein hints in training and prediction [26]. Comparison of protein hints generated by the two pipelines would not be straightforward since BRAKER2 uses hints to introns and start/stop codons while MAKER2 uses hints to exon parts. More accurate training of AUGUSTUS was supposed to be one of the important factors for elevated accuracy of BRAKER2. A simple and effective way to improve MAKER2 accuracy was to reduce the number of gene finders from three to two (Fig. S5, Tables 4, S9, S10).

The difference in accuracy of MAKER2 and BRAKER2 could be even larger for eukaryotic genomes with longer length, however, such a comparison is more difficult to make due to less accurate annotations. Therefore, since a comprehensive comparison of the two methods is not a goal of this paper, the presented results are limited to the three well studied genomes.

Last but not least, the training of gene finders is not fully automated in MAKER2. Even though there are recommended training protocols, users still have to execute the training steps manually. On the other hand, BRAKER2 can be executed with a single command, from start to finish.

### Comparison with BRAKER1

We saw that gene prediction accuracy of BRAKER2 depended on the volume of reference proteins. Similarly, the accuracy of BRAKER1 depends on the volume of the RNA-Seq data. For running BRAKER1 on genomes of *A. thaliana, C. elegans* and *D. melanogaster*, we sampled collections of RNA-Seq from SRA that covered the three genomes very well. We observed that the accuracy of BRAKER2 was consistently comparable or better than the one of BRAKER1 when we used the largest set of the proteins for each species, the relevant OrthoDB partition exempting proteins of the same species.

Both BRAKER1 and BRAKER2 predicted a rather low percentage of annotated alternative isoforms in genomes of *A. thaliana, C. elegans* and *D. melanogaster* (Table S14). This result is explained by the parameter setting in AUGUSTUS that ignored an RNA-Seq or protein hint if there was a contradicting hint supported 10 times more frequently. The reference genome annotations of the three species are rather inclusive in a sense of collecting lowly expressed isoforms.

### EuGene and other gene finders

A gene finder for eukaryotic genomes, EuGene, provides a mechanism for the integration of several sources of information into the gene prediction process. This modular tool can integrate data derived from protein spliced alignment to genomic DNA [47]. Unfortunately, EuGene does not provide recommendations on model training for the case when protein sequences are the only source of the external evidence [47]. For this reason, we have not used this tool in this comparative study.

Several tools attempt accurate identification of gene structures in a novel genome by mapping homologous proteins (e.g. GenomeThreader [28], Scipio [30]). This approach limits the gene discovery to genes of homologs present in the input protein set; the accuracy of this method drops significantly with increase of evolutionary distance between the two species [10, 45]. Another significant challenge for a number of earlier developed tools is the processing of large volumes of proteins. This challenge is much smaller if a tool, like GeMoMa [48, 49], is oriented on getting the protein information from closely related species. In addition, GeMoMa requires gene coordinates in reference genomes. We did not make comparisons of BRAKER2 to the above mentioned tools since these tools were not designed for the situation when the reference protein data does not contain proteins from closely related species.

## Conclusions

BRAKER2 is a fully automated tool for gene prediction in a novel eukaryotic genome. It allows to leverage information accumulated in protein databases. BRAKER2 generates hints from millions of proteins in the course of several hours (for instance, in case of *D. melanogaster*, ∼2.6 millions of proteins were processed in ∼3 hours). In the tests on genomes of plants, insects and other animals, we observed that BRAKER2 delivered state-of-the-art annotation accuracy and was favorably compared to already existing tools.

## AVAILABILITY

BRAKER2 is available at https://github.com/Gaius-Augustus/BRAKER. All additional scripts and data used to generate figures and tables in this manuscript are available at https://github.com/gatech-genemark/BRAKER2-exp.

## Link to Supplement

https://docs.google.com/document/d/1aBbbqMpIEfNwi-b6Py44cVYocyoUqCqWgUgbbpqeQRg/edit?usp=sharing

