## Supplemental Materials for "BRAKER2: Automatic Eukaryotic Genome Annotation with GeneMark-EP+ and AUGUSTUS Supported by a Protein Database"

#### Supplementary Figures

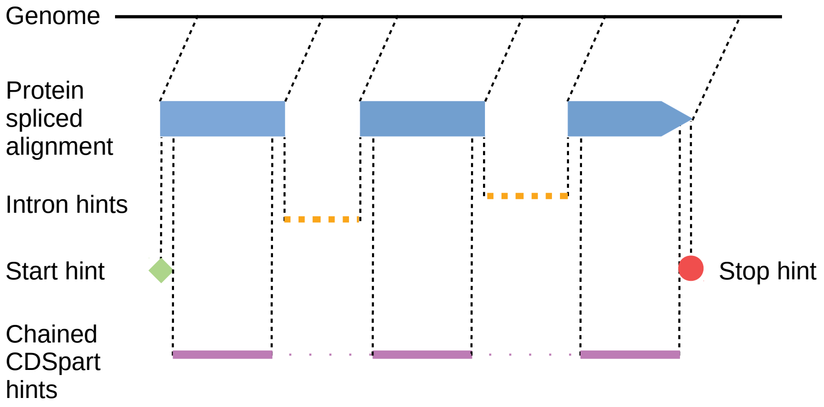


Figure S1: Schematic illustration of hints derived by ProtHint from a spliced alignment of a protein to genomic sequence. GeneMark-EP+ and AUGUSTUS treat intron hints as independent ones, not necessarily related to one and the same gene. CDSpart hints indicate locations of protein-coding exons (CDS). Each CDSpart hint is trimmed at its boundaries by 15 nucleotides. CDSpart hints originating from the same protein are treated as a 'chain' of evidence, i.e. all such CDSpart hints are incorporated into the same transcript model.

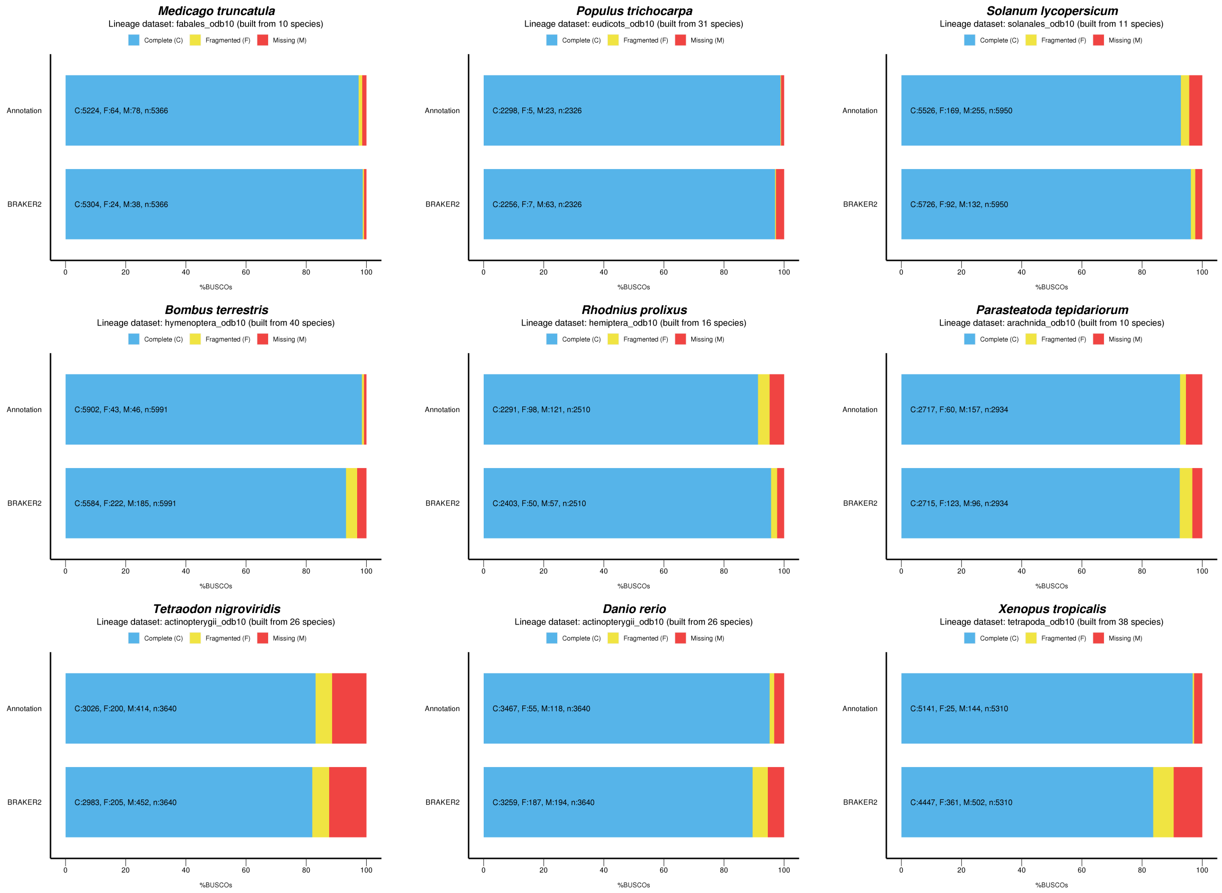


Figure S2: Estimation of a completeness of the set of genes from BUSCO families present in the reference genome annotation (top in each panel) and predicted by BRAKER2 (bottom in each panel)

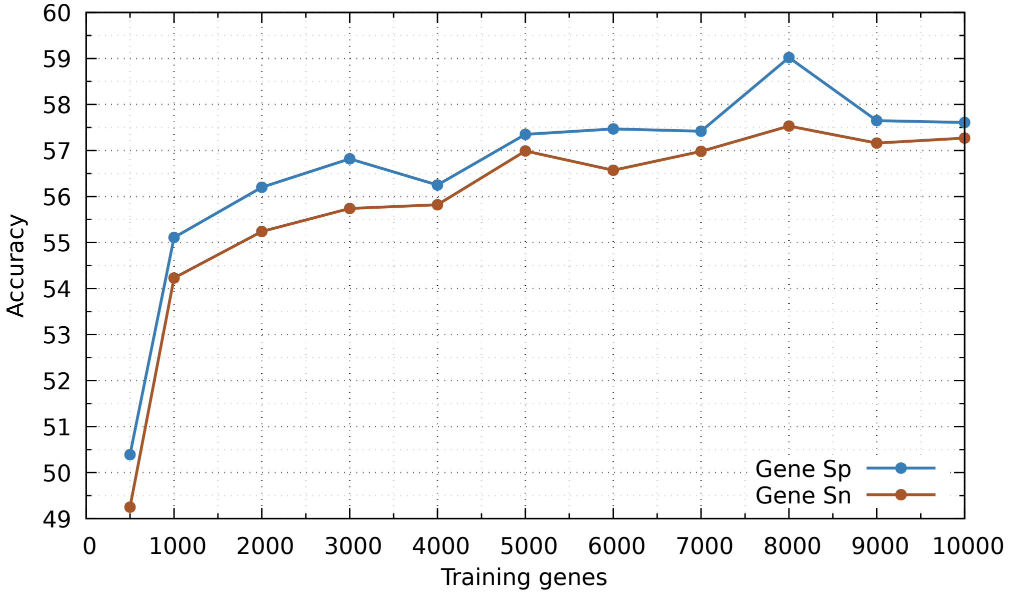


Figure S3: Dependence of Sn and Sp of the AUGUSTUS *ab initio* gene prediction in the *A. thaliana* genome on the number of anchored genes in training. When the number of training genes was very large, AUGUSTUS training became a time-consuming step. Therefore, we experimented with a number of anchored genes used for training in *A. thaliana* since BRAKER2 generates more than 10,000 anchored genes for this species. For training AUGUSTUS, we used 500, 1,000, 2,000, ..., 10,000 anchored genes selected at random. A very low Sn and Sp were observed for 500 genes. Both Sn and Sp jumped upon increase of the number of genes to 1,000 and then increased almost steadily when the number of genes increased from 1,000 to 8,000 genes. While we selected 8,000 as the upper limit for the number of training genes, we consider 4,000 as the minimum necessary for training. In a practical setting, if the number of all fully anchored genes is less than 4,000 for a given genome, more genes are added to the training set by BRAKER2 in order of the level of support by the protein hints (Supplementary Materials, Section 1.1). In the experiments related to this figure the gene sets larger than 8,000 were not reduced. The supporting proteins from the Plantae segment of OrthoDB did not include proteins from the *A. thaliana* genus.

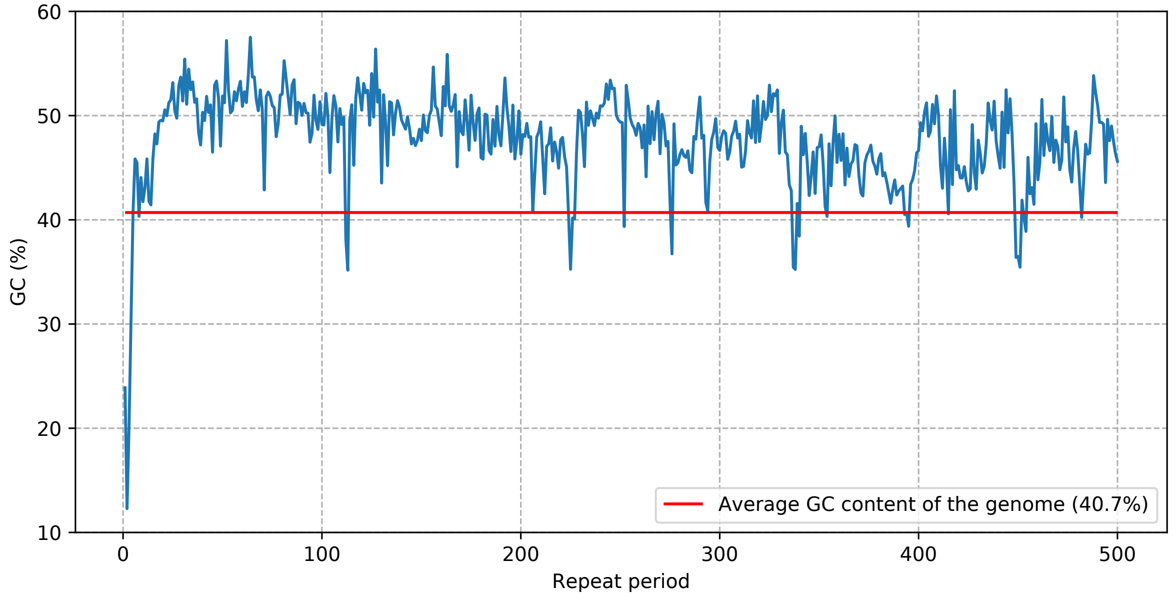


Figure S4: GC-content of tandem repeats in the *X. tropicalis* genome shown as a function of the size of repeat period.

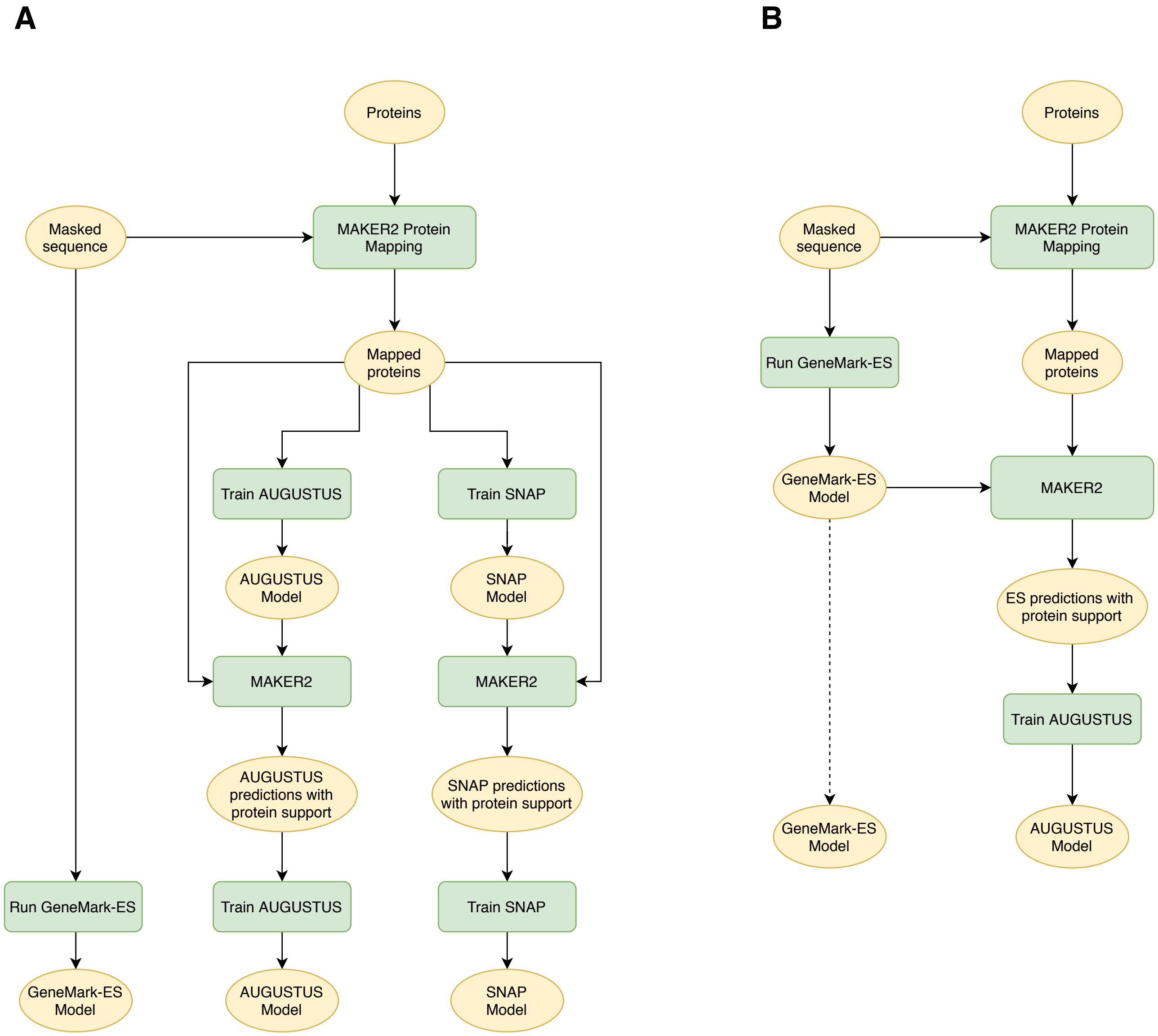


Figure S5: Schematics of the MAKER2 training protocols: (A) a protocol recommended by the MAKER2 authors [1] (B) a new protocol (similar to one of BRAKER2) that was implemented and produced better gene prediction accuracy.

#### Supplementary Tables

| Species | Assembly version |
|---|---|
| Species with early sequenced genomes | |
| *Arabidopsis thaliana* | GCF_000001735 |
| *Caenorhabditis elegans* | GCA_001483305 |
| *Drosophila melanogaster* | GCA_000001215 |
| Other species | |
| Plantae | |
| *Populus trichocarpa* | Ptrichocarpa_533_v4.0 |
| *Medicago truncatula* | GCA_003473485.2 |
| *Solanum lycopersicum* | SL4.0 |
| Arthropoda | |
| *Bombus terrestris* | GCF_000214255.1 |
| *Rhodnius prolixus* | GCA_000181055.3 |
| *Parasteatoda tepidariorum* | GCF_000365465.2 |
| Vertebrata | |
| *Tetraodon nigroviridis* | TETRAODON 8.0 |
| *Danio rerio* | GCF_000002035 |
| *Xenopus tropicalis* | GCF_000004195.4 |

Table S1: Genomes assembly versions used for testing BRAKER2.

| Species | Gene | | | Exon | | | % Non-canonical or incomplete genes |
|---|---|---|---|---|---|---|---|
| | Sn | Sp | F1 | Sn | Sp | F1 | |
| *P. trichocarpa* | 69.1 | 60.2 | 64.3 | 84.9 | 82.3 | 83.6 | 0.3 |
| *M. truncatula* | 44.7 | 44.0 | 44.3 | 78.7 | 71.5 | 74.9 | 0.0 |
| *S. lycopersicum* | 41.2 | 34.4 | 37.5 | 76.6 | 67.7 | 71.9 | 14.5 |
| *B. terrestris* | 46.9 | 25.0 | 32.6 | 74.5 | 72.0 | 73.2 | 4.7 |
| *R. prolixus* | 16.0 | 10.6 | 12.8 | 60.6 | 49.7 | 54.6 | 34.7 |
| *P. tepidariorum* | 30.4 | 14.9 | 20.0 | 67.7 | 59.6 | 63.4 | 18.2 |
| *T. nigroviridis* | 11.0 | 7.9 | 9.2 | 60.5 | 56.7 | 58.5 | 63.8 |
| *D. rerio* | 40.6 | 20.5 | 27.2 | 75.3 | 69.4 | 72.2 | 11.8 |
| *X. tropicalis* | 40.6 | 25.9 | 31.6 | 75.1 | 77.5 | 76.3 | 2.4 |

Table S2: Complementary information for Table 3 in the main text; Sn, Sp and F1 values computed on exon and gene level. For a gene to be considered complete and canonical, at least one of the gene's transcripts had to be fully annotated, with initial exon starting with a 'canonical' ATG and a terminal exon ending with TAA, TAG or TGA.

| *A. thaliana* | *C. elegans* | *D. melanogaster* |
|---|---|---|
| *Dendrobium officinale* | *Buceros rhinoceros silvestris* | *Pogonomyrmex barbatus* |
| *Parasponia andersonii* | *Cardiocondyla obscurior* | *Oryctes borbonicus* |
| *Beta vulgaris subsp. vulgaris* | *Drosophila elegans* | *Heliconius melpomene* |
| *Aegilops tauschii* | *Geospiza fortis* | *Stegodyphus mimosarum* |
| *Nelumbo nucifera* | *Sarcoptes scabiei* | *Calopteryx splendens* |
| *Triticum urartu* | *Austrofundulus limnaeus* | *Wasmannia auropunctata* |
| *Ananas comosus* | *Nomascus leucogenys* | *Fopius arisanus* |
| *Coccomyxa subellipsoidea C-169* | *Pieris rapae* | *Limulus polyphemus* |
| *Populus euphratica* | *Anas platyrhynchos* | *Tribolium castaneum* |
| *Phalaenopsis equestris* | *Numida meleagris* | *Myzus cerasi* |

Table S3: Proteins of these species were used as external evidence in tests of MAKER2 and BRAKER2. The three groups of ten species were selected at random from the OrthoDB partitions (see main text).

| | BRAKER2 | | | |
|---|---|---|---|---|
| | Order excluded | Family excluded | Species excluded | BRAKER1 |
| Gene Sn | 71.1 | 73.6 | 79.4 | 61.6 |
| Gene Sp | 67.0 | 69.7 | 72.8 | 61.7 |
| Gene F1 | 69.0 | 71.6 | 76.0 | 61.6 |
| Exon Sn | 80.7 | 81.5 | 83.3 | 79.9 |
| Exon Sp | 86.6 | 87.4 | 86.8 | 81.7 |
| Exon F1 | 83.5 | 84.3 | 85.0 | 80.8 |

Table S4: Gene prediction accuracy of BRAKER2 and BRAKER1 observed in tests on the *A. thaliana* genome. The sets of reference proteins for BRAKER2 were selected from the Plantae section of OrthoDB.

| | BRAKER2 | | | |
|---|---|---|---|---|
| | Order excluded | Family excluded | Species excluded | BRAKER1 |
| Gene Sn | 49.8 | 49.1 | 67.4 | 58.2 |
| Gene Sp | 56.2 | 55.1 | 68.3 | 62.3 |
| Gene F1 | 52.8 | 51.9 | 67.8 | 60.2 |
| Exon Sn | 75.4 | 74.7 | 84.3 | 83.6 |
| Exon Sp | 88.6 | 88.2 | 90.7 | 87.2 |
| Exon F1 | 81.5 | 80.9 | 87.4 | 85.4 |

Table S5: The same type of information as in Table S4 for a test on the *C. elegans* genome. The sets of reference proteins for BRAKER2 were selected from the Metazoa section of OrthoDB.

| | BRAKER2 | | | |
|---|---|---|---|---|
| | Order excluded | Family excluded | Species excluded | BRAKER1 |
| Gene Sn | 61.1 | 66.3 | 77.8 | 63.1 |
| Gene Sp | 60.2 | 64.8 | 72.9 | 61.8 |
| Gene F1 | 60.6 | 65.5 | 75.3 | 62.4 |
| Exon Sn | 71.4 | 74.5 | 79.8 | 76.7 |
| Exon Sp | 83.2 | 85.1 | 87.6 | 80.7 |
| Exon F1 | 76.8 | 79.4 | 83.5 | 78.6 |

Table S6: The same type of information as in Table S4 for a test on the *D. melanogaster* genome. The sets of reference proteins for BRAKER2 were selected from the Arthropoda section of OrthoDB.

| | *A. thaliana* | | *C. elegans* | | *D. melanogaster* | |
|---|---|---|---|---|---|---|
| | All proteins | Subset 10 | All proteins | Subset 10 | All proteins | Subset 10 |
| Intron Sn | 69.4 | 68.5 | 18.2 | 10.1 | 33.8 | 29.8 |
| Intron Sp | 98.8 | 99.2 | 99.3 | 99.4 | 99.0 | 99.6 |
| Start Sn | 36.3 | 27.8 | 5.5 | 3.3 | 15.6 | 11.1 |
| Stop Sp | 94.7 | 95.0 | 95.5 | 97.1 | 94.6 | 96.3 |
| Start Sn | 34.9 | 29.2 | 8.1 | 5.1 | 19.4 | 14.7 |
| Stop Sp | 97.7 | 98.2 | 97.4 | 98.7 | 99.0 | 99.2 |

Table S7: Accuracy of ProtHint high-confidence hints to introns and gene border sites. We show results of experiments for complete set of reference proteins from relevant OrthoDB partitions (exempting species of the same taxonomic order) or for proteins from 10 randomly selected species from the same OrthoDB partitions (Table S3).

| | *C. elegans* | | *A. thaliana* | | *D. melanogaster* | | *D. rerio* | |
|---|---|---|---|---|---|---|---|---|
| | All | Anchored* | All | Anchored | All | Anchored | All | Anchored |
| Gene Sn | 38.6 | **43.7** | 52.8 | **55.9** | 51.7 | **54.4** | 15.0 | **26.5** |
| Gene Sp | 46.2 | **50.9** | 55.5 | **56.9** | 52.2 | **55.7** | 7.3 | **13.1** |
| Gene F1 | 42.1 | **47.0** | 54.1 | **56.4** | 51.9 | **55.0** | 9.8 | **17.5** |
| Exon Sn | **75.5** | 75.1 | **75.9** | 75.7 | 68.5 | **68.6** | 68.3 | **73.5** |
| Exon Sp | 83.4 | **86.2** | 81.6 | **83.2** | 76.2 | **80.5** | 50.4 | **63.0** |
| Exon F1 | 79.3 | **80.3** | 78.6 | **79.3** | 72.1 | **74.1** | 58.0 | **67.8** |

Table S8: *Ab initio* prediction accuracy of AUGUSTUS trained on i/ *All* genes predicted by GeneMark-EP+ and ii/ *Anchored* genes (see Methods in the main text). The results for the first three species were generated with reference proteins from species outside a taxonomic family of a relevant species, for *D. rerio* we used proteins from species outside of taxonomic order. (*) When $<$ 4000 anchored genes were available, additional genes were added in the descending order of their support by protein hints to reach 4000 genes (see Supplementary Methods, Section 1.1 for details). This approach was used for *C. elegans* that had 2,332 anchored genes.

| Species | *A. thaliana* | | | *C. elegans* | | | *D. melanogaster* | | |
|---|---|---|---|---|---|---|---|---|---|
| Training | MAKER2 / BRAKER2-like | | | MAKER2 / BRAKER2-like | | | MAKER2 / BRAKER2-like | | |
| Predictors | SNAP GM-ES AUGUSTUS | GM-ES AUGUSTUS | AUGUSTUS | SNAP GM-ES AUGUSTUS | GM-ES AUGUSTUS | AUGUSTUS | SNAP GM-ES AUGUSTUS | GM-ES AUGUSTUS | AUGUSTUS |
| Gene Sn | 49.3 / 50.6 | 52.9 / **53.9** | 48.5 / 49.8 | 25.5 / 26.2 | 28.4 / **30.4** | 24.6 / 26.6 | 42.6 / 44.6 | 45.0 / **48.0** | 42.8 / 46.2 |
| Gene Sp | 42.1 / 43.8 | 54.1 / **55.5** | 49.9 / 51.8 | 22.1 / 23.0 | 37.1 / **38.9** | 32.1 / 34.0 | 31.1 / 31.5 | 46.8 / **50.3** | 44.8 / 48.8 |
| Gene F1 | 45.4 / 47.0 | 53.5 / **54.7** | 49.2 / 50.8 | 23.6 / 24.5 | 32.2 / **34.1** | 27.9 / 29.8 | 35.9 / 37.0 | 45.9 / **49.2** | 43.8 / 47.5 |
| Exon Sn | 73.4 / 73.8 | 74.5 / **74.7** | 72.5 / 72.7 | 61.7 / **63.8** | 59.7 / 62.6 | 58.3 / 61.2 | 62.8 / **64.3** | 61.7 / 63.7 | 60.4 / 62.5 |
| Exon Sp | 72.6 / 72.9 | **83.4** / 83.0 | 82.1 / 81.5 | 64.5 / 65.0 | 80.6 / **81.4** | 78.3 / 79.2 | 58.7 / 54.6 | 75.3 / **76.0** | 74.3 / 75.1 |
| Exon F1 | 73.0 / 73.3 | **78.7** / 78.6 | 77.0 / 76.8 | 63.1 / 64.4 | 68.6 / **70.8** | 66.9 / 69.0 | 60.7 / 59.1 | 67.8 / **69.3** | 66.6 / 68.2 |

Table S9: Prediction accuracy of MAKER2 on repeat-masked genomes. The table shows results generated by using gene finders trained directly on gene structures derived by protein alignments (MAKER2), as recommended by MAKER2 authors; and by using gene prediction tools trained on genes predicted by GeneMark-ES and supported at least partially by protein alignments (BRAKER2-like). Three combinations of gene finders in MAKER2 (SNAP + GeneMark-ES + AUGUSTUS; GeneMark-ES + AUGUSTUS; AUGUSTUS) are compared.

| Species | *A. thaliana* | | | *C. elegans* | | | *D. melanogaster* | | |
|---|---|---|---|---|---|---|---|---|---|
| Training | MAKER2 / BRAKER2-like | | | MAKER2 / BRAKER2-like | | | MAKER2 / BRAKER2-like | | |
| Predictors | SNAP GM-ES AUGUSTUS | GM-ES AUGUSTUS | AUGUSTUS | SNAP GM-ES AUGUSTUS | GM-ES AUGUSTUS | AUGUSTUS | SNAP GM-ES AUGUSTUS | GM-ES AUGUSTUS | AUGUSTUS |
| Gene Sn | 52.9 / 54.3 | 58.6 / **59.2** | 53.5 / 54.7 | 34.9 / 34.5 | 43.0 / **43.9** | 36.0 / 39.3 | 46.1 / 48.3 | 50.1 / **52.0** | 45.8 / 49.3 |
| Gene Sp | 35.1 / 36.7 | 45.4 / 46.7 | 46.2 / **49.0** | 25.8 / 25.5 | 38.1 / 38.9 | 43.7 / **46.8** | 24.1 / 26.0 | 35.8 / 37.8 | 42.0 / **45.8** |
| Gene F1 | 42.2 / 43.8 | 51.2 / **52.2** | 49.6 / 51.7 | 29.7 / 29.3 | 40.4 / 41.3 | 39.5 / **42.8** | 31.7 / 33.8 | 41.8 / 43.8 | 43.8 / **47.5** |
| Exon Sn | 75.7 / 76.0 | **77.6** / 77.4 | 75.2 / 75.1 | 75.9 / 76.6 | 78.2 / **79.2** | 69.2 / 72.5 | 65.7 / **67.1** | 65.5 / 66.8 | 62.2 / 64.5 |
| Exon Sp | 62.8 / 63.3 | 72.3 / 72.1 | **76.6** / 75.7 | 66.0 / 64.8 | 77.7 / 78.1 | 84.2 / **85.2** | 46.5 / 46.2 | 60.0 / 60.8 | 69.8 / **69.7** |
| Exon F1 | 68.7 / 69.1 | 74.8 / 74.7 | **75.9** / 75.4 | 70.6 / 70.2 | 77.9 / **78.7** | 76.0 / 78.3 | 54.4 / 54.7 | 62.6 / 63.7 | 65.8 / **67.0** |

Table S10: Same comparison as in Table S9, with gene predictions made on unmasked genomes.

| | *A. thaliana* | | | | *C. elegans* | | | | *D. melanogaster* | | | |
|---|---|---|---|---|---|---|---|---|---|---|---|---|
| | BRAKER2 | | | BUSCO | BRAKER2 | | | BUSCO | BRAKER2 | | | BUSCO |
| | Order excl. | Family excl. | Species excl. | | Order excl. | Family excl. | Species excl. | | Order excl. | Family excl. | Species excl. | |
| Gene Sn | 52.0 | 55.9 | **57.3** | 47.7 | 45.2 | 43.7 | **49.1** | 24.2 | 54.2 | 54.4 | **55.3** | 51.4 |
| Gene Sp | 52.9 | 56.9 | **58.4** | 55.0 | 52.9 | 50.9 | **55.0** | 35.9 | 55.0 | 55.7 | 55.2 | **57.5** |
| Gene F1 | 52.4 | 56.4 | **57.8** | 51.1 | 48.7 | 47.0 | **51.9** | 28.9 | 54.6 | 55.0 | **55.2** | 54.3 |
| Exon Sn | 74.1 | 75.7 | **76.8** | 73.0 | 76.3 | 75.1 | **78.3** | 62.3 | 68.2 | 68.6 | **69.2** | 66.2 |
| Exon Sp | 81.2 | 83.2 | 83.7 | **84.3** | 86.9 | 86.2 | **87.4** | 80.1 | 79.9 | 80.5 | 79.0 | **82.0** |
| Exon F1 | 77.5 | 79.3 | **80.1** | 78.2 | 81.3 | 80.3 | **82.6** | 70.1 | 73.6 | **74.1** | 73.8 | 73.3 |

Table S11: *Ab initio* gene prediction accuracy of AUGUSTUS with model parameters estimated on the training set generated by BRAKER2 and with model parameters estimated on a training set made from genes predicted by AUGUSTUS-PPX [2] with evidence from BUSCO families processed via the BUSCO protocol (see Supplementary Results, Section 2.2).

**A**

| *A. thaliana* — Family excluded proteins | BRAKER2 steps 1 and 2: GeneMark ES | GeneMark EP+ | BRAKER2 steps 3 and 4: AUGUSTUS with hints, first iteration | second iteration |
|---|---|---|---|---|
| Gene Sn | 55.8 | 67.5 | 73.2 | 73.6 |
| Gene Sp | 54.0 | 64.6 | 69.4 | 69.7 |
| Gene F1 | 54.9 | 66.0 | 71.3 | 71.6 |
| Exon Sn | 77.2 | 80.3 | 81.3 | 81.5 |
| Exon Sp | 79.2 | 83.7 | 87.3 | 87.4 |
| Exon F1 | 78.2 | 81.9 | 84.2 | 84.3 |

| *C. elegans* — Family excluded proteins | BRAKER2 steps 1 and 2: GeneMark ES | GeneMark EP+ | BRAKER2 steps 3 and 4: AUGUSTUS with hints, first iteration | second iteration |
|---|---|---|---|---|
| Gene Sn | 46.8 | 47.4 | 48.9 | 49.1 |
| Gene Sp | 46.4 | 45.8 | 54.9 | 55.1 |
| Gene F1 | 46.6 | 46.6 | 51.7 | 51.9 |
| Exon Sn | 81.0 | 80.3 | 74.7 | 74.7 |
| Exon Sp | 82.4 | 81.5 | 88.1 | 88.2 |
| Exon F1 | 81.7 | 80.9 | 80.8 | 80.9 |

| *D. melanogaster* — Family excluded proteins | BRAKER2 steps 1 and 2: GeneMark ES | GeneMark EP+ | BRAKER2 steps 3 and 4: AUGUSTUS with hints, first iteration | second iteration |
|---|---|---|---|---|
| Gene Sn | 50.2 | 59.5 | 65.6 | 66.3 |
| Gene Sp | 47.6 | 56.1 | 64.1 | 64.8 |
| Gene F1 | 48.9 | 57.7 | 64.8 | 65.6 |
| Exon Sn | 67.6 | 71.9 | 74.2 | 74.5 |
| Exon Sp | 72.0 | 78.2 | 84.9 | 85.1 |
| Exon F1 | 69.7 | 74.9 | 79.2 | 79.5 |

**B**

| *A. thaliana* — Species excluded proteins | BRAKER2 steps 1 and 2: GeneMark ES | GeneMark EP+ | BRAKER2 steps 3 and 4: AUGUSTUS with hints, first iteration | second iteration |
|---|---|---|---|---|
| Gene Sn | 55.8 | 73.7 | 78.9 | 79.4 |
| Gene Sp | 54.0 | 69.4 | 72.7 | 72.9 |
| Gene F1 | 54.9 | 71.5 | 75.7 | 76.0 |
| Exon Sn | 77.2 | 81.8 | 83.1 | 83.3 |
| Exon Sp | 79.2 | 84.8 | 86.8 | 86.7 |
| Exon F1 | 78.2 | 83.2 | 84.9 | 85.0 |

| *C. elegans* — Species excluded proteins | BRAKER2 steps 1 and 2: GeneMark ES | GeneMark EP+ | BRAKER2 steps 3 and 4: AUGUSTUS with hints, first iteration | second iteration |
|---|---|---|---|---|
| Gene Sn | 46.8 | 53.4 | 66.8 | 67.4 |
| Gene Sp | 46.4 | 51.8 | 67.7 | 68.3 |
| Gene F1 | 46.6 | 52.6 | 67.2 | 67.8 |
| Exon Sn | 81.0 | 82.4 | 84.1 | 84.3 |
| Exon Sp | 82.4 | 84.1 | 90.6 | 90.7 |
| Exon F1 | 81.7 | 83.2 | 87.2 | 87.4 |

| *D. melanogaster* — Species excluded proteins | BRAKER2 steps 1 and 2: GeneMark ES | GeneMark EP+ | BRAKER2 steps 3 and 4: AUGUSTUS with hints, first iteration | second iteration |
|---|---|---|---|---|
| Gene Sn | 50.2 | 69.2 | 76.6 | 77.8 |
| Gene Sp | 47.6 | 63.1 | 72.0 | 72.9 |
| Gene F1 | 48.9 | 66.0 | 74.2 | 75.3 |
| Exon Sn | 67.6 | 76.2 | 79.3 | 79.8 |
| Exon Sp | 72.0 | 80.9 | 87.3 | 87.6 |
| Exon F1 | 69.7 | 78.5 | 83.1 | 83.5 |

Table S12: Change of the gene prediction accuracy upon successive execution of components of BRAKER2, on the *three* genomes with reference proteins from the relevant OrthoDB partitions with A/ proteins from the same taxonomic family excluded, and B/ proteins from the same species excluded.

| C. elegans | Order excluded proteins | | | | Family excluded proteins | | | | Species excluded proteins | | | |
|---|---|---|---|---|---|---|---|---|---|---|---|---|
| | HC Hints | HC Hints LC Hints | HC Hints Chains | HC Hints LC Hints Chains | HC Hints | HC Hints LC Hints | HC Hints Chains | HC Hints LC Hints Chains | HC Hints | HC Hints LC Hints | HC Hints Chains | HC Hints LC Hints Chains |
| Gene Sn | 47.5 | 49.6 | 49.3 | 49.8 | 45.9 | 48.8 | 48.5 | 49.1 | 55.1 | 65.9 | 65.1 | 67.4 |
| Gene Sp | 54.9 | 56.6 | 55.9 | 56.2 | 52.8 | 55.2 | 54.6 | 55.1 | 58.7 | 67.1 | 66.4 | 68.3 |
| Gene F1 | 50.9 | 52.9 | 52.4 | 52.8 | 49.1 | 51.8 | 51.4 | 51.9 | 56.9 | 66.5 | 65.7 | 67.8 |
| Exon Sn | 73.8 | 75.1 | 75.2 | 75.4 | 72.5 | 74.1 | 74.3 | 74.7 | 78.7 | 83.4 | 83.2 | 84.3 |
| Exon Sp | 88.7 | 89.0 | 88.7 | 88.6 | 88.0 | 88.5 | 88.3 | 88.2 | 89.4 | 91.0 | 90.9 | 90.7 |
| Exon F1 | 80.6 | 81.5 | 81.4 | 81.5 | 79.5 | 80.7 | 80.7 | 80.9 | 83.7 | 87.0 | 86.9 | 87.4 |

| A.thaliana | Order excluded proteins | | | | Family excluded proteins | | | | Species excluded proteins | | | |
|---|---|---|---|---|---|---|---|---|---|---|---|---|
| | HC Hints | HC Hints LC Hints | HC Hints Chains | HC Hints LC Hints Chains | HC Hints | HC Hints LC Hints | HC Hints Chains | HC Hints LC Hints Chains | HC Hints | HC Hints LC Hints | HC Hints Chains | HC Hints LC Hints Chains |
| Gene Sn | 65.0 | 68.5 | 70.2 | 71.1 | 66.9 | 70.7 | 73.3 | 73.6 | 72.9 | 76.7 | 79.1 | 79.4 |
| Gene Sp | 63.6 | 66.0 | 66.4 | 67.0 | 65.8 | 68.5 | 69.6 | 69.7 | 69.6 | 71.6 | 73.2 | 72.9 |
| Gene F1 | 64.3 | 67.2 | 68.3 | 69.0 | 66.3 | 69.6 | 71.4 | 71.6 | 71.2 | 74.1 | 76.0 | 76.0 |
| Exon Sn | 78.5 | 79.9 | 80.4 | 80.7 | 79.1 | 80.6 | 81.2 | 81.5 | 81.3 | 82.6 | 83.1 | 83.3 |
| Exon Sp | 86.3 | 86.6 | 86.6 | 86.6 | 87.0 | 87.4 | 87.6 | 87.4 | 86.8 | 86.6 | 87.2 | 86.7 |
| Exon F1 | 82.2 | 83.1 | 83.4 | 83.6 | 82.9 | 83.8 | 84.3 | 84.3 | 83.9 | 84.5 | 85.1 | 85.0 |

| D. melanogaster | Order excluded proteins | | | | Family excluded proteins | | | | Species excluded proteins | | | |
|---|---|---|---|---|---|---|---|---|---|---|---|---|
| | HC Hints | HC Hints LC Hints | HC Hints Chains | HC Hints LC Hints Chains | HC Hints | HC Hints LC Hints | HC Hints Chains | HC Hints LC Hints Chains | HC Hints | HC Hints LC Hints | HC Hints Chains | HC Hints LC Hints Chains |
| Gene Sn | 58.6 | 60.2 | 60.4 | 61.1 | 62.8 | 64.6 | 65.7 | 66.3 | 73.7 | 76.0 | 77.5 | 77.8 |
| Gene Sp | 59.0 | 60.1 | 59.8 | 60.2 | 63.1 | 63.9 | 64.7 | 64.8 | 71.0 | 72.1 | 73.0 | 72.9 |
| Gene F1 | 58.8 | 60.1 | 60.1 | 60.6 | 63.0 | 64.3 | 65.2 | 65.6 | 72.3 | 74.0 | 75.2 | 75.3 |
| Exon Sn | 69.4 | 70.5 | 70.9 | 71.4 | 72.2 | 73.5 | 74.0 | 74.5 | 78.0 | 79.1 | 79.5 | 79.8 |
| Exon Sp | 83.3 | 83.5 | 83.3 | 83.2 | 84.9 | 85.0 | 85.4 | 85.1 | 87.2 | 87.4 | 87.9 | 87.6 |
| Exon F1 | 75.7 | 76.5 | 76.6 | 76.8 | 78.1 | 78.8 | 79.3 | 79.5 | 82.3 | 83.0 | 83.5 | 83.5 |

Table S13: Accuracy of BRAKER2 determined for various combinations of the types of external evidence generated from spliced aligned proteins. HC and LC stand for high and low confidence, respectively.

| | | *A. thaliana* | | | |
|---|---|---|---|---|---|
| | | Genes | Transcripts | Alternative transcripts | % Alt from all |
| | Annotation | 27,444 | 40,827 | 13,383 | 32.8 |
| | BRAKER1 | 27,403 | 28,899 | 1,496 | 5.2 |
| BRAKER2 with exclusion of proteins from | Species | 29,902 | 31,844 | 1,942 | 6.1 |
| | Family | 28,988 | 30,153 | 1,165 | 3.9 |
| | Order | 29,101 | 30,248 | 1,147 | 3.8 |

| | | *C. elegans* | | | |
|---|---|---|---|---|---|
| | | Genes | Transcripts | Alternative transcripts | % Alt from all |
| | Annotation | 20,172 | 28,506 | 8,334 | 29.2 |
| | BRAKER1 | 18,833 | 20,978 | 2,145 | 10.2 |
| BRAKER2 with exclusion of proteins from | Species | 19,916 | 21,366 | 1,450 | 6.8 |
| | Family | 17,977 | 18,466 | 489 | 2.6 |
| | Order | 17,883 | 18,283 | 400 | 2.2 |

| | | *D. melanogaster* | | | |
|---|---|---|---|---|---|
| | | Genes | Transcripts | Alternative transcripts | % Alt from all |
| | Annotation | 13,929 | 22,247 | 8,318 | 37.4 |
| | BRAKER1 | 14,208 | 15,470 | 1,262 | 8.2 |
| BRAKER2 with exclusion of proteins from | Species | 14,863 | 16,149 | 1,286 | 8.0 |
| | Family | 14,247 | 15,266 | 1,019 | 6.7 |
| | Order | 14,142 | 14,605 | 463 | 3.2 |

Table S14: Numbers of genes and transcripts predicted by BRAKER1 and BRAKER2 with various sets of reference proteins (proteins from the same species, family and order excluded from the OrthoDB partitions).

### 1 Methods

#### 1.0 Definitions of Sensitivity (Sn), Specificity (Sp) and Harmonic Mean (F1)

Let us consider a set $S$ of objects possessing one of two properties (designated by signs plus and minus). Let us consider a property prediction method $M$ applied with the goal to identify all the objects that have the 'plus' property, a subset $S$+. Therefore, the method may not be necessarily applied to all the objects in $S$ individually. The method $M$ is characterized by the numbers 'True positives' ($TP$) – number of correctly identified 'positive' objects, 'False negatives' ($FN$) – number of positive objects identified as 'negative', and 'False positives' ($FP$) – number of negative objects, identified as positive.

For method $M$, the sensitivity value (Sn) is defined as $TP/(TP + FN)$, the specificity value (Sp) is defined as $TP/(TP + FP)$, and the harmonic mean of Sn and Sp, $(2 \times \text{Sn} \times \text{Sn})/(\text{Sn} + \text{Sp})$, is designated as F1.

#### 1.1 Selection of GeneMark-EP+ predicted genes for training AUGUSTUS

Genes predicted by GeneMark-EP+ [3] are filtered and sampled prior training AUGUSTUS [4,5] in the following way:

1. The ratio of multi-exon and single-exon genes is determined prior to filtering.
2. During filtering, multi-exon genes are retained if they have support by an intron hint from at least one protein alignment in every exon.
3. The minimal number of required single-exon genes in relation to filtered multi-exon genes is computed to keep the proportion from step 1 in step 4.
4. Single-exon genes are selected if they have support from protein evidence in terms of start- and stop-codon hints. If the number of the selected single-exon genes is lower than the minimal required number of single-exon genes, then the single-exon genes predicted by GeneMark-EP+ that do not have protein evidence support are randomly selected until the minimal number is reached.
5. If the resulting number of training genes is lower than 4000, additional genes are added in the diminishing order of their support rank by protein hints. A gene support rank is computed as follows:

   $$S_r = \frac{\#of\ supported\ borders\ of\ protein\text{-}coding\ exons}{\#of\ actual\ borders\ of\ protein\text{-}coding\ exons}$$

   Genes are then added in the descending order of their $S_r$.
6. Complex genes with many introns contribute more effectively to training AUGUSTUS than gene structures with few or no introns. Such simpler organized genes are therefore down-sampled as described earlier [6].
7. Training genes are translated into protein sequences that are searched against each other. If two sequences have an identity of more than 80%, one gene is removed from the training gene set.
8. If there are more than 8000 training gene structures, genes are randomly down-sampled to 8000 genes to decrease runtime of BRAKER2.

Within BRAKER2, the training gene set for AUGUSTUS is randomly split into three sets:

1. A set for running *etraining*, the tool for training AUGUSTUS parameters,
2. a set for evaluating parameter optimization steps (within `optimize_augustus.pl`), and
3. a test set. This set is used as an independent test set for estimating the accuracy.

If the total number of genes is smaller than 600, 1/3 of all available genes will be sampled into each set. If there are 600 to 1000 available gene structures, 200 genes each are sampled into the last two sets, all remaining genes go into the first set. If there are more than 1000 training gene structures, 300 genes each are sampled into the last two sets, all remaining genes go into the first set.

#### 1.2 Classification of intron hints

For gene prediction with AUGUSTUS, intron hints that do not fall into the category of high confidence hints are further separated into medium and low confidence hints. For this, BRAKER2 uses logistic regression with parameters that were obtained with ProtHint [3] using the *Drosophila melanogaster* genome and Arthropoda section of OrthoDB [7]. ProtHint intron hints were labeled as true or false using the reference annotation. A hold-out test-set of 500 hints was set aside. The binomial logistic regression model was made to predict whether a hint was true or false using multiplicity and the alignment score of ProtHint with R as follows:

```
glm.fit <- glm(label ~ mult_norm + al_score, data = trainset, family=binomial(link='logit'))
```

Accuracy of the model was checked on the test set:

```
fitted.results <- predict(glm.fit ,newdata=testset, type='response')
fitted.results <- ifelse(fitted.results > 0.5,1,0)

misClasificError <- mean(fitted.results != testset$label)
print(paste('Accuracy',1-misClasificError))
```

Accuracy was 93% (the proportion of true positives in the test set was 80%).

The resulting coefficients are used by BRAKER2 to classify intron hints from 'non high confidence' class of ProtHint:

```
            Estimate Std. Error z value Pr(>|z|)
(Intercept) -4.00529    0.04935  -81.16   <2e-16 ***
mult_norm    4.73909    0.04662  101.66   <2e-16 ***
al_score     9.09026    0.14741   61.67   <2e-16 ***
```

We confirmed that these parameters work reasonably well on *Arabidopsis thaliana*.

#### 1.3 Extrinsic evidence configuration parameters in AUGUSTUS in BRAKER2

Extrinsic parameters for evidence integration with AUGUSTUS were adapted using *Arabidopsis thaliana* genome and hints generated with ProtHint using OrthoDB v10 Plants section (exempting proteins from the same species). Final non-neutral extrinsic parameters used for all species by BRAKER2 were:

```
[SOURCES]
M RM P C

# M: manual hints, to be enforced hints
# RM: repeats
# P: protein hints
# C: chained protein hints

[GENERAL]
start       1     1      M 1 1e+100 RM 1    1 P 2 1 1e3 1e6 C 1  1e6
stop        1     1      M 1 1e+100 RM 1    1 P 2 1 1e3 1e6 C 1  1e6
ass         1     1    1 M 1 1e+100 RM 1    1 P 2 1 1e2 1e2 C 1  1e2
dss         1     1    1 M 1 1e+100 RM 1    1 P 2 1 1e2 1e2 C 1  1e2
intron      1 0.168      M 1 1e+100 RM 1    1 P 2 1 1e2 100 C 1 3.16
CDSpart     1     1 0.99 M 1 1e+100 RM 1    1 P 2 1 1e2 1e4 C 1  1e4
nonexonpart 1     1      M 1 1e+100 RM 1 1.14 P 2 1   1   1 C 1    1
```

#### 1.4 Preparation of input data

##### 1.4.1 Genomes

Table S1 shows which assembly version was used for each species. The assemblies were processed with commands documented at `https://github.com/gatech-genemark/EukSpecies-BRAKER2/tree/master/`. The processing steps included, for instance, renaming chromosomes/contigs to match the names in annotation as well as removal of mitochondrial and chloroplast DNA.

##### 1.4.2 Repeat Masking

Genomes were *de novo* masked for repeats with a combination of RepeatModeler [8] (open-1.0.11) and RepeatMasker [9] (1.332) with the following commands:

```
BuildDatabase -engine wublast -name genome genome.fasta
RepeatModeler -engine wublast -database genome
RepeatMasker  -engine wublast -lib genome-families.fa -xsmall genome.fasta
```

Additional masking by Tandem Repeats Finder [10] (with maximum repeat period size = 500) was applied to *X. tropicalis* since the default run of RepeatMasker/RepeatModeler did not identify a significant portion of long tandem repeats (with repeat pattern length > 10) in this genome.

Tandem Repeats Finder (TRF) v4.07b was run with the following command:

```
trf genome.fasta 2 7 7 80 10 50 500 -d -m -h
```

We then converted the coordinates of repeats from TRF `.dat` format to `.gff` with a custom `parseTrfOutput.py`[1] script:

```
parseTrfOutput.py genome.fasta.2.7.7.80.10.50.500.dat --minCopies 1 --statistics STATS \
    --gc > genome.fasta.2.7.7.80.10.50.500.raw.gff
```

Next, we soft-masked the genome using these coordinates:

```
# Sort gff
sort -k1,1 -k4,4n -k5,5n genome.fasta.2.7.7.80.10.50.500.raw.gff > sorted
# Merge overlapping repeats
bedtools merge -i sorted | awk 'BEGIN{OFS="\t"} \
    {print $1,"trf","repeat",$2+1,$3,".",".",".","."}' \
    > genome.fasta.2.7.7.80.10.50.500.merged.gff
# Apply masking
bedtools maskfasta -fi genome.fasta.masked -bed \
    genome.fasta.2.7.7.80.10.50.500.merged.gff \
    -fo genome.fasta.combined.masked -soft
```

In the above commands, `genome.fasta.masked` is the genome masked by RepeatMasker/RepeatModeler and `genome.fasta.combined.masked` is the final masking combining RepeatMasker/RepeatModeler and additional TRF masking.

Figure S4, describing GC content per repeat period size, was generated with the `plot_stats.py`[2] script:

```
plot_stats.py STATS.GC xt-gc.pdf --title "X. tropicalis: GC content per \
    repeat period size" --scaleY 0.01 --line 40.7 --ymin 10 --ymax 60
```

**Reproducing the Masking** Different runs of masking by RepeatModeler/RepeatMasker may result in slightly different masking coordinates due to the stochasticity of the algorithms. For this reason, we uploaded the masking coordinates used in the BRAKER2 project at `https://github.com/gatech-genemark/EukSpecies-BRAKER2/tree/master/${SPECIES}/annot/mask.gff.gz`. These coordinates make it possible to soft-mask the genome without having to re-run RepeatModeler and RepeatMasker. To soft mask the genome with the coordinates, use the following commands:

```
gunzip mask.gff.gz
bedtools maskfasta -fi genome.fasta -bed mask.gff -fo genome.fasta.masked -soft
```

The final masking coordinates for *X. tropicalis* are in the same repository, in a file named `combined.mask.gff.gz`.

[1] https://github.com/gatech-genemark/BRAKER2-exp/blob/master/bin/trf-scripts/parseTrfOutput.py
[2] https://github.com/gatech-genemark/BRAKER2-exp/blob/master/bin/trf-scripts/plot_stats.py

##### 1.4.3 Protein Databases

For each species, we used OrthoDB v10 [7] proteins from a corresponding taxonomic phylum or kingdom (see Table 2 in the main text). From this protein set, we excluded proteins of species from the same taxonomic order (resulting in `order_excluded.fa` file) to simulate the absence of proteins of closely related species. For each of the three organisms (*C. elegans*, *A. thaliana*, and *D. melanogaster*), we also prepared larger protein sets by excluding proteins from the same taxonomic family and proteins of the species of interest itself (`family_excluded.fa` and `species_excluded.fa` files).

For example, the proteins for the case of *D. melanogaster* were prepared in the following way:

```
# Download arthropoda proteins from OrthoDB

wget https://v100.orthodb.org/download/odb10_arthropoda_fasta.tar.gz
tar xvf odb10_arthropoda_fasta.tar.gz
rm odb10_arthropoda_fasta.tar.gz

# Function for creating a single fasta file with arthropoda proteins,
# excluding species supplied in a list.

createProteinFile() {
  excluded=$1
  output=$2

  # Get NCBI ids of species in excluded list
  grep -f <(paste <(yes $'\n'| head -n $(cat $excluded | wc -l)) \
  $excluded <(yes $'\n'| head -n $(cat $excluded | wc -l))) \
  ../../OrthoDB/odb10v0_species.tab | cut -f2 > ids

  # Create protein file with everything else
  cat $(ls -d arthropoda/Rawdata/* | grep -v -f ids) > $output

  # Remove dots from file
  sed -i -E "s/\.//" $output

  rm ids
}

# Create protein databases with different levels of exclusion.
# Exclusion lists correspond to species in taxonomic levels in OrthoDB v10.

createProteinFile drosophila_melanogaster.txt species_excluded.fa
createProteinFile drosophila.txt family_excluded.fa
createProteinFile diptera.txt order_excluded.fa
```

Here, `diptera.txt`, `drosophila.txt` and `drosophila_melanogaster.txt` contain the lists of names of species in the same taxonomic order, family, and *D. melanogaster* itself.

A Description of protein preparation for each of the tested species is available at `https://github.com/gatech-genemark/BRAKER2-exp/tree/master/${SPECIES}/data`.

#### 1.5 Running BRAKER2

BRAKER2 version (2.1.6) with GeneMark-EP+ (4.58), AUGUSTUS (3.3.4), and ProtHint (2.5.0) was run with the following command:

```
braker.pl --genome genome.fasta.masked --prot_seq order_excluded.fasta --softmasking
```

For each of the three organisms (*C. elegans*, *A. thaliana*, and *D. melanogaster*), we also ran BRAKER2 with `species_excluded.fasta` and `family_excluded.fasta` proteins (See 1.4.3).

##### 1.5.1 Running BRAKER1

BRAKER1 [11] was run with the following command:

```
braker.pl --genome genome.fasta.masked --hints varus.gff --softmasking
```

See section 1.10 for details about RNA-Seq sampling and mapping.

#### 1.6 Selection of genes for training

Table S8 was generated as follows. To get *ab initio* predictions of AUGUSTUS trained on anchored genes from outcome of GeneMark-EP+, we ran BRAKER2 with default settings (see 1.5) along with an additional `--AUGUSTUS_ab_initio` option.

To get *ab initio* predictions of AUGUSTUS trained on all GeneMark-EP+ genes, we used a modified BRAKER2 version 2.1.6. To reproduce the results, replace `braker.pl` with `braker_es_all.pl`[3] and run the following command:

```
braker_es_all.pl --genome genome.fasta.masked --geneMarkGtf genemark_ep.gtf \
    --softmasking --esmode --AUGUSTUS_ab_initio
```

In the above command, `genemark_ep.gtf` was taken from the standard BRAKER2 run. The accuracy was computed (see 1.11) for the `augustus.ab_initio.gtf` results.

#### 1.7 Number of training genes

Figure S3 was generated as follows. First, we extended `braker.pl` with an option `--maxTrainGenes` to control the maximum number of training genes in BRAKER2. To use this modified version, get BRAKER2 version 2.1.6 and replace braker.pl with `braker_max_train_genes.pl`[4] .

Then, we ran BRAKER2 with different limits on the number of training genes:

```
echo "$(echo 500; seq 1000 1000 10000)" | xargs -I {} braker_max_train_genes.pl \
    --maxTrainGenes {} --genome genome.fasta.masked --softmasking \
    --skipIterativePrediction --geneMarkGtf genemark_ep.gtf --hints hintsfile.gff \
    --prothints prothint.gff --evidence evidence.gff --AUGUSTUS_ab_initio \
    --epmode --workingdir max_{}
```

The files `genemark_ep.gtf`, `hintsfile.gff`, `prothint.gff`, and `evidence.gtf` were taken from a standard BRAKER2 run.

At the next step, we computed AUGUSTUS *ab initio* accuracy (see 1.11 for details about accuracy computation). This result is available in the `ab_initio.genes`[5] file.

Finally, we used the `max_train_genes_accuracies.gp`[6] Gnuplot script to generate the Figure S3:

```
gnuplot -e "in='ab_initio.genes'" max_train_genes_accuracies.gp
```

[3]https://github.com/gatech-genemark/BRAKER2-exp/blob/master/bin/braker_es_all.pl
[4]https://github.com/gatech-genemark/BRAKER2-exp/blob/master/bin/max_train_genes/braker_max_train_genes.pl
[5]https://github.com/gatech-genemark/BRAKER2-exp/blob/master/bin/max_train_genes/ab_initio.genes
[6]https://github.com/gatech-genemark/BRAKER2-exp/blob/master/bin/max_train_genes/max_train_genes_accuracies.gp

#### 1.8 AUGUSTUS training on the BUSCO genes

We trained AUGUSTUS using a gene set predicted by AUGUSTUS-PPX within BUSCO Software [2, 12, 13] for *A. thaliana* with *eudicotyledons_odb10* and *tomato* as starting species, for *C. elegans* with *metazoa_odb9* and *schistosoma* as starting species, and for *D. melanogaster* with *arthropoda_odb9* and *ant* as starting species. Parameters trained on these 'BUSCO genes' were used by BRAKER2 for predicting genes with AUGUSTUS in *ab initio* mode (skipping training of AUGUSTUS, see section 1.8.1).

BUSCO 3.0.2 was used to train AUGUSTUS 3.3.3 in the long time execution mode (including a run of `optimize_augustus.pl`). The commands are shown below:

```
# on Drosophila melanogaster:
python3 run_BUSCO.py -i genome.fasta.masked -o busco_arthropoda_ant \
    -l arthropoda_odb9 -m geno -c 11 -sp ant --long
# on Arabidopsis thaliana:
python3 run_BUSCO.py -i genome.fasta.masked -o busco_eudicotyledons_tomato \
    -l eudicotyledons_odb10 -m geno -c 11 -sp tomato --long
# on Caenorhabditis elegans:
python3 run_BUSCO.py -i genome.fasta.masked -o busco_metazoa_schistosoma \
    -l metazoa_odb9 -m geno -c 11 -sp schistosoma --long
```

The *ant* parameters for the run on *Drosophila melanogaster* have been generated using BRAKER1 on a yet unpublished high quality genome assembly. The parameter set is available on request and will be included in the next AUGUSTUS release after publication of that genome.

##### 1.8.1 AUGUSTUS *ab initio* gene predictions with the BUSCO gene derived parameters

AUGUSTUS was run in *ab initio* mode using parameters trained on BUSCO set for each genome (parameter sets *BUSCO_busco_arthropoda_ant_3655623124*,
*BUSCO_busco_eudicotyledons_tomato_2767133850*, and
*BUSCO_busco_metazoa_schistosoma_807538653*
are available at `https://github.com/Gaius-Augustus/Augustus/tree/master/config/species`):

```
braker.pl --species ${BUSCO_PARAMS} --softmasking --esmode \
    --skipAllTraining --genome genome.fasta.masked
```

#### 1.9 Experiments with MAKER2

MAKER2 (version 3.01.03, MPI mode) [14] was run with the gene finders SNAP (release from 11/29/2013) [15], AUGUSTUS (version 3.3.3) [5], and GeneMark-ES [16] (version 4.58).

We ran MAKER2 in two distinct ways (Figure S5): (i) using a protocol recommended for novel species by the MAKER2 authors [1] and (ii) a protocol based on training procedure developed for BRAKER2.

Final predictions were generated with MAKER2 using several alternative gene finder combinations: (i) SNAP, GeneMark-ES, and AUGUSTUS, (ii) GeneMark-ES and AUGUSTUS, (iii) AUGUSTUS only.

All training steps of MAKER2 were executed on a repeat-masked sequence. Final predictions were run on both masked and unmasked sequences.

To reduce runtime of MAKER2, ten species from a relevant OrthoDB partition were randomly selected for each of the three model organisms (Table S3). This selection procedure is described at `https://github.com/gatech-genemark/BRAKER2-exp/tree/master/${SPECIES}/data`. BRAKER2 was run with the same subsets of species.

The comparison of accuracy of ProtHint with all OrthoDB partition species and just the subset of ten is shown in Table S7.

##### 1.9.1 Repeat masking in MAKER2 (difference with BRAKER2)

For repeat masking, MAKER2 used the same repeat libraries (generated by RepeatModeler) as BRAKER2 (for details, see section 1.4.2 about repeat masking).

It is important to note that even though MAKER2 and BRAKER2 used the same repeat libraries, genomes are masked differently by MAKER2 when comparing to our approach of using RepeatMasker/RepeatModeler for BRAKER2.

MAKER2 runs RepeatMasker internally. Subsequently, MAKER2 does hard-masking of all interspersed (complex) repeats while low-complexity (simple) repeats remain soft-masked. Borders of complex repeats are extended by 50 nt.

For the BRAKER2 run, the sequence was soft-masked with RepeatMasker. Within BRAKER2, AUGUSTUS uses information on soft-masked regions at the gene prediction step and reduces the probability of coding exon prediction in repeat regions. GeneMark-ES and -EP+ do hard-masking of soft-masked repeats longer than 1000 nt for genomes shorter than 300 Mb and they do hard-masking of soft-masked repeats longer than 100 nt for genomes longer than 300 Mb.

##### 1.9.2 Running MAKER2 with a protocol recommended by the authors

For running MAKER2 in a way recommended by authors of MAKER2 for novel species, we followed the tutorials at `http://weatherby.genetics.utah.edu/MAKER/wiki/index.php/MAKER_Tutorial_for_GMOD_Online_Training_2014`, `https://weatherby.genetics.utah.edu/MAKER/wiki/index.php/MAKER_Tutorial_for_WGS_Assembly_and_Annotation_Winter_School_2018`, and [1], mostly. Running MAKER2 in this mode consists of seven steps:

1. Training GeneMark-ES,
2. generating training gene structures for SNAP and AUGUSTUS on the basis of protein to genome alignment with MAKER2,
3. training SNAP and AUGUSTUS on initial training genes,
4. predicting genes with MAKER2 using initial parameters of SNAP and protein sequences to produce better training genes for SNAP,
5. predicting genes with MAKER2 using initial parameters of AUGUSTUS and protein sequences to produce better training genes for AUGUSTUS,
6. retraining SNAP and AUGUSTUS on genes predicted in steps 4. and 5., respectively,
7. predicting genes with MAKER2 using GeneMark-ES, SNAP, AUGUSTUS, and protein sequences.

Compiling training genes and training AUGUSTUS and SNAP requires numerous manual steps because MAKER2 is not an automated pipeline for training any gene finder. The description of training steps is documented in sections 1.9.2.1 – 1.9.2.7.

###### 1.9.2.1 Training GeneMark-ES

GeneMark-ES was trained using the command

```
cd /${SPECIES}/ES

gmes_petap.pl --soft_mask auto --ES genome.fasta.masked
```

###### 1.9.2.2 Generation of training genes for SNAP and AUGUSTUS Iteration 1

MAKER2 configuration files were generated:

```
cd /${SPECIES}/protein_mapping

maker -CTL
```

The file maker_opts.ctl was edited to contain (besides default parameters):

```
genome=genome.fasta
protein=proteins.fa
protein2genome=1
model_org=
rmlib=genome-families.fa
repeat_protein=
```

No HMMs for gene finders were configured. MAKER2 was run with the following command:

```
mpiexec maker maker_opts.ctl maker_bopts.ctl maker_exe.ctl
```

Predicted genes and aligned protein hints were collected with the commands:

```
# gff3_merge belongs to MAKER2
gff3_merge -d genome.maker.output/genome_master_datastore_index.log
cat genome.maker.output/genome_datastore/*/*/*/*/evidence*.gff > evidence.gff
```

###### 1.9.2.3 Training SNAP Iteration 1

MAKER2 predictions were converted to ann/zff and dna files (native format for training SNAP) and SNAP was subsequently trained:

```
cd /${SPECIES}/snap_protein_training_1

# maker2zff belongs to MAKER2
maker2zff -n ../protein_mapping/genome.all.gff

# fathom, forge and hmm-assembler.pl are part of SNAP
fathom -categorize 1000 genome.ann genome.dna
fathom -export 1000 -plus uni.ann uni.dna
forge export.ann export.dna
hmm-assembler.pl ${SPECIES} . > ${SPECIES}.hmm
```

###### 1.9.2.4 Training AUGUSTUS Iteration 1

To obtain a training gene file for AUGUSTUS, the training set in "ann" format already generated for SNAP was first converted to gff3, and from there to gtf format:

```
cd /${SPECIES}/augustus_protein_training_1

# zff2gff3.pl belongs to SNAP
zff2gff3.pl ../snap_protein_training_1/genome.ann > all.gff3

cat all.gff3 | perl -ne '
  if(not(m/^\#/)){
    chomp; @t = split(/\t/);
```

```
    @t2 = split(/=/, $t[7]);
    print "$t[0]\t$t[1]\t$t[2]\t$t[3]\t$t[4]\t$t[5]\t$t[6]\t";
    print "\tgene_id \"$t2[1]_$t[0]\"; transcript_id";
    print " \"$t2[1]_$t[0]\"\n";
  }
' > all.gtf
```

The flanking region for AUGUSTUS training genes was computed comparable to BRAKER2:

```
cat all.gtf | perl -ne '
  @t = split(/\t/);
  $seen{$t[8]} += ($t[4] - $t[3] + 1);
  if(eof()){
    $sum = 0; $c = 0;
    foreach my $key ( keys %seen ){
      $c=$c+1; $sum += $seen{$key};
    }
    print ($sum."/".$c."=".($sum/$c))/2;
    print "\n";
  }
'
```

The flanking region length `$F_LENGTH` (result of the previous command) was used when excising training genes from the genome.

```
# gff2gbSmallDNA.pl belongs to AUGUSTUS
gff2gbSmallDNA.pl all.gtf genome.fasta.masked $F_LENGTH first.gb
```

Note that in BRAKER2, in practice, the actual flanking region is frequently shorter than the computed flanking region value, which is only an upper boundary in any case, because GeneMark-EP+ predicts genes in *ab initio* mode that limits flanking region size. In this MAKER2 protocol, the flanking region is much less often limited by a neighboring gene because only evidence derived genes go into the training set.

AUGUSTUS was trained as follows:

```
# new_species.pl is part of AUGUSTUS
new_species.pl --species=${SPECIES}_from_proteins
```

The original training gene structures contained incomplete genes (missing start- or stop-codons). Such genes were filtered out:

```
# etraining and filgerGenesOut_mRNAname.pl are part of AUGUSTUS
etraining --species=${SPECIES}_from_proteins first.gb 1> etrain-test.out 2> etrain-test.err
fgrep "gene" etrain-test.err | cut -f 2 -d " " > bad.etraining-test.lst
filterGenesOut_mRNAname.pl bad.etraining-test.lst first.gb > second.gb
```

In case of *A. thaliana*, more than 8000 training genes remained in `second.gb`. Such a large number increases runtime of training AUGUSTUS but usually does not lead to a large increase in accuracy. Therefore, we randomly selected 8000 genes to proceed with training (same threshold as used in BRAKER2):

```
mv second.gb second.gb.fullset
randomSplit.pl second.gb.fullset 8000
mv second.gb.fullset.test second.gb
```

Here, we proceed with the file `second.gb` for all species. The training gene set was split into two sets, the second set was subsequently further split in another two sets, resulting in three different files:

1. A small test set of 300 genes for measuring accuracy after etraining and optimize_augustus.pl,

2. a large gene set for etraining, that was further split into:
    (a) a large gene set for for the option `--onlytrain` of optimize_augustus.pl,
    (b) a small gene set for optimize_augustus.pl, the size was 300.

```
# randomSplit.pl is part of AUGUSTUS
randomSplit.pl second.gb 300
randomSplit.pl second.gb.train 300
# this results in the following files:
# 1) second.gb.test -> measuring accuracy
# 2) second.gb.train -> etraining
# 2a) second.gb.train.train -> --onlytrain in optimize_augustus.pl
# 2b) second.gb.train.test -> optimize_augustus.pl
```

Major AUGUSTUS parameters were adjusted with *etraining*:

```
etraining --species=${SPECIES}_from_proteins second.gb.train
# modify stop codon frequencies manually
```

*etraining* does not automatically modify stop codon frequencies in the model files. The stop codon frequencies, in a file

`/path/to/augustus/config/species/${SPECIES}_from_proteins/${SPECIES}_from_proteins_parameters.cfg`

were modified manually based on *etraining* output.

Other parameters were optimized with optimize_augustus.pl:

```
optimize_augustus.pl --species=${SPECIES}_from_proteins --onlytrain=second.gb.train.train \
    second.gb.train.test
```

###### 1.9.2.5 Generating training genes for SNAP Iteration 2

MAKER2 parameters were generated:

```
cd /${SPECIES}/maker_snap

maker -CTL
```

The file maker_opts.ctl was edited to contain (besides default parameters):

```
genome=genome.fasta
protein=
protein_gff=../protein_mapping/evidence.gff
model_org=
rmlib=genome-families.fa
repeat_protein=
snaphmm=../snap_protein_training_1/${SPECIES}.hmm
```

Notice that the protein fasta file (option `protein=`) was replaced with aligned hints generated in the previous iteration of MAKER2 (`protein_gff=../protein_mapping/evidence.gff`). This significantly speeds up the computation because protein mapping is one of the most time consuming parts of the MAKER2 pipeline.

Another way to speed up the computation is to copy the folder with intermediate files from the previous MAKER2 run. MAKER2 is then able to re-use parts which do not change between the different runs.

```
cp -r ../protein_mapping/genome.maker.output .
```

MAKER2 was run with the following command:

```
mpiexec maker maker_opts.ctl maker_bopts.ctl maker_exe.ctl
gff3_merge -d genome.maker.output/genome_master_datastore_index.log
```

###### 1.9.2.6 Generating training genes for AUGUSTUS Iteration 2

MAKER2 parameters were generated:

```
cd /${SPECIES}/maker_augustus

maker -CTL
```

The file maker_opts.ctl was edited to contain (besides default parameters):

```
genome=genome.fasta
protein=
protein_gff=../protein_mapping/evidence.gff
model_org=
rmlib=genome-families.fa
repeat_protein=
augustus_species=${SPECIES}_from_proteins
```

Intermediate files were copied from previous MAKER run (further details in section 1.9.2.5)

```
cp -r ../protein_mapping/genome.maker.output .
```

MAKER2 was run with the following command:

```
mpiexec maker maker_opts.ctl maker_bopts.ctl maker_exe.ctl
gff3_merge -d genome.maker.output/genome_master_datastore_index.log
```

###### 1.9.2.7 Retraining SNAP and AUGUSTUS

**SNAP** Retraining of SNAP was done in the same way as described in section 1.9.2.3. The following lines:

```
cd /${SPECIES}/snap_protein_training_1
maker2zff -n ../protein_mapping/genome.all.gff
```

were replaced with:

```
cd /${SPECIES}/snap_protein_training_2
maker2zff -n ../maker_snap/genome.all.gff
```

**AUGUSTUS** Retraining of AUGUSTUS was done in the same way as described in section 1.9.2.4. The following lines:

```
cd /${SPECIES}/augustus_protein_training_1
zff2gff3.pl ../snap_protein_training_1/genome.ann > all.gff3
```

were replaced with:

```
cd /${SPECIES}/augustus_protein_training_2
maker2zff -n ../maker_augustus/genome.all.gff
zff2gff3.pl genome.ann > all.gff3
```

Additionally, no new species was created in the second round of AUGUSTUS training, meaning that the following command was not used:

```
new_species.pl --species=${SPECIES}_from_proteins
```

###### 1.9.2.8 Predicting genes with MAKER2

MAKER2 parameters were generated:

```
cd /${SPECIES}/maker_final_prediction

maker -CTL
```

The file maker_opts.ctl was edited to contain (besides default parameters):

```
genome=genome.fasta
protein=
protein_gff=../protein_mapping/evidence.gff
model_org=
rmlib=genome-families.fa
repeat_protein=
augustus_species=${SPECIES}_from_proteins
snaphmm=../snap_protein_training_2/${SPECIES}.hmm
gmhmm=../ES/gmhmm.mod
keep_preds=1
```

Intermediate files were copied from the previous MAKER run (further details in section 1.9.2.5)

```
cp -r ../protein_mapping/genome.maker.output .
```

MAKER2 was run with the following command:

```
mpiexec maker maker_opts.ctl maker_bopts.ctl maker_exe.ctl
gff3_merge -d genome.maker.output/genome_master_datastore_index.log
```

Predictions were converted to .gtf format with GenomeTools [17]:

```
gt gff3_to_gtf <(grep -P "^#|\tmaker\t" genome.all.gff) > maker.gtf
```

The MAKER2 prediction accuracy was evaluated as described in section 1.11.

###### 1.9.2.9 Additional gene prediction modes

To run MAKER without any of the three gene predictors, we simply left one (or more) of the following options empty:

```
augustus_species=${SPECIES}_from_proteins
snaphmm=../snap_protein_training_2/${SPECIES}.hmm
gmhmm=../ES/gmhmm.mod
```

For MAKER runs on an unmasked sequence, we left the `rmlib` option empty:

```
rmlib=
```

##### 1.9.3 BRAKER2-like MAKER2 protocol

As an alternative to the recommended MAKER2 training, we tested a protocol in which AUGUSTUS and SNAP are trained on the basis of protein-supported GeneMark-ES predictions. This protocol is thus more similar to the way training is executed within BRAKER2. Running MAKER2 in this mode consists of four steps:

1. Training GeneMark-ES,
2. predicting genes with MAKER2 using GeneMark-ES and protein sequences,
3. training SNAP and AUGUSTUS on protein-supported genes predicted in step 2.,
4. predicting genes with MAKER2 using GeneMark-ES, SNAP, AUGUSTUS, and protein sequences.

###### 1.9.3.1 Training GeneMark-ES

GeneMark-ES was trained in the same way as described in section 1.9.2.1, using the command

```
cd /${SPECIES}/ES
gmes_petap.pl --soft_mask auto --ES genome.fasta.masked
```

###### 1.9.3.2 Generation of training sets for SNAP and AUGUSTUS

MAKER2 configuration files were generated:

```
cd /${SPECIES}/masker_es
maker -CTL
```

The file maker_opts.ctl was edited to contain (besides default parameters):

```
genome=genome.fasta
protein=proteins.fa
gmhmm=../ES/gmhmm.mod
model_org=
rmlib=genome-families.fa
repeat_protein=
```

The rest of the code was run in the same way as described in Section 1.9.2.2.

###### 1.9.3.3 Training SNAP and AUGUSTUS

**SNAP** SNAP was trained in the same way as described in section 1.9.2.3. The following lines:

```
cd /${SPECIES}/snap_protein_training_1
maker2zff -n ../protein_mapping/genome.all.gff
```

were replaced with:

```
cd /${SPECIES}/snap_es_training
maker2zff -n ../maker_es/genome.all.gff
```

**AUGUSTUS** Training of AUGUSTUS was done in the same way as described in section 1.9.2.4. The following lines:

```
cd /${SPECIES}/augustus_protein_training_1
zff2gff3.pl ../snap_protein_training_1/genome.ann > all.gff3
```

were replaced with:

```
cd /${SPECIES}/augustus_es_training
maker2zff -n ../maker_es/genome.all.gff
zff2gff3.pl genome.ann > all.gff3
```

Additionally, the AUGUSTUS species name (`--species=${SPECIES}_from_proteins` flag) was changed to `--species=${SPECIES}_from_es`.

###### 1.9.3.4 Predicting genes with MAKER2

MAKER2 parameters were generated:

```
cd /${SPECIES}/maker_final_prediction_braker_like

maker -CTL
```

The file maker_opts.ctl was edited to contain (besides default parameters):

```
genome=genome.fasta
protein=
protein_gff=../protein_mapping/evidence.gff
model_org=
rmlib=genome-families.fa
repeat_protein=
augustus_species=${SPECIES}_from_es
snaphmm=../snap_es_training/${SPECIES}.hmm
gmhmm=../ES/gmhmm.mod
keep_preds=1
```

The rest of the code was run in the same way as described in Section 1.9.2.8.

##### 1.9.4 Running BRAKER2 for comparison with MAKER2

BRAKER2 was executed with the same protein sets as MAKER2 as follows:

```
braker.pl --genome=genome.fasta.masked --prot_seq=proteins.fa --softmasking --cores=8
```

#### 1.10 Running VARUS to sample and align RNA-Seq libraries

VARUS [18] (version from March 26, 2020) with fastq-dump [19] (v2.10.4) and HISAT2 [20] (v2.1.0) was run with the following command:

```
runVARUS.pl --aligner=HISAT --readFromTable=0 --createindex=1 --latinGenus=$GENUS \
    --latinSpecies=$SPECIES --speciesGenome=genome.fasta.masked
```

Stranded introns mapped by VARUS are located in `$GENUS_$SPECIES/cumintrons.stranded.gff`, this result is referred to as `varus.gff` in this document.

Results of VARUS depend on the date VARUS was run since the amount of data deposited to NCBI Sequence Read Archive [19], from which VARUS samples reads, is changing in time. Therefore, we uploaded the result of VARUS for each species here `https://github.com/tomasbruna/braker2-exp/tree/master/${SPECIES}/varus`. The aforementioned folder also contains information on when VARUS was run and what specific VARUS parameters (`VARUSparameters.txt`) were used.

#### 1.11 Accuracy evaluation

##### 1.11.1 Annotation parsing

Table 1 in the main text shows which annotation version was used for each species. These annotations were processed to generate a uniform annotation format. The processing steps (documented at `https://github.com/gatech-genemark/EukSpecies-BRAKER2/${SPECIES}`) are composed of, for example, categorization of complete and incomplete genes or partition of pseudogenic regions into a separate file. The processed annotations are available in the same repository in the `annot` folder.

##### 1.11.2 Evaluation against a full set of annotated genes

Prediction accuracy against the whole set of annotated genes (Tables S4, S5, and S6) was automatically computed by BRAKER2 with `compute_accuracies.sh`[7]. To run `compute_accuracies.sh` separately (for gene,

[7] `https://github.com/Gaius-Augustus/BRAKER/blob/master/scripts/compute_accuracies.sh`

transcript, and exon levels), the following command can be used:

```
compute_accuracies.sh annot.gtf annot_pseudo.gff prediction.gtf gene trans cds
```

Annotated pseudogenic regions were excluded from the accuracy computation, i.e. predicted pseudogenes do not count as false positives.

###### 1.11.3 Accuracy in Figs. 3 and 4

Figs. 3 and 4 in the main text were generated with the `visualize_distances_results.sh`[8] script:

```
visualize_distances_results.sh annot.gtf outputFolder {gene,cds} [xmin xmax ymin ymax]
```

###### 1.11.4 Evaluation of accuracy against RNA-Seq supported gene sets

Prediction sensitivity of BRAKER2 computed against a subset of annotated complete multi-exon genes that have all introns supported by at least one RNA-Seq read sampled by VARUS (see 1.10), shown in Table 3 in the main text, was calculated with the `complete_supported_subset_table.sh`[9] script in the following way:

```
complete_supported_subset_table.sh prediction.gtf completeTranscripts.gtf \
    annot_pseudo.gff varus.gff
```

###### 1.11.5 Plots depicting completeness of the sets of predicted genes in the BUSCO families

Each of the BUSCO [12, 13] completeness plots (Figure S2) was generated with BUSCO (v4.0.5) for genes in annotation and BRAKER2 predictions with the following commands:

```
busco -m protein -i augustus.hints.aa -o BRAKER2 -l ${LINEAGE}
busco -m protein -i annot.aa -o ANNOT -l ${LINEAGE}

mkdir plot
cp BRAKER2/short_summary*.txt plot
cp ANNOT/short_summary*.txt plot

python3 generate_plot_unified_completeness.py -wd plot
```

In the commands above, `${LINEAGE}` is the name of a BUSCO lineage dataset. Lineage datasets used for each species are displayed in the Figure S2. The file `augustus.hints.aa` contains BRAKER2 protein predictions and it is a part of the standard BRAKER2 output. File `annot.aa` contains proteins translated from annotation. We used the `getAnnoFastaFromJoingenes.py`[10] script to translate the annotated genes:

```
getAnnoFastaFromJoingenes.py -g genome.fasta.masked -o annot -f annot.gtf
```

To generate the BUSCO figures, we used the `generate_plot_unified_completeness.py`[11] script, which is a modification of an original BUSCO script available at `https://gitlab.com/ezlab/busco/-/blob/master/scripts/generate_plot.py`.

[8] https://github.com/gatech-genemark/BRAKER2-exp/blob/master/bin/visualize_distances_results.sh
[9] https://github.com/gatech-genemark/BRAKER2-exp/blob/master/bin/complete_supported_subset_table.sh
[10] https://github.com/Gaius-Augustus/Augustus/blob/master/scripts/getAnnoFastaFromJoingenes.py
[11] https://github.com/gatech-genemark/BRAKER2-exp/blob/master/bin/generate_plot_unified_completeness.py

### 2 Results

#### 2.1 MAKER2 predictions and accuracy

We ran MAKER2 [14] in two distinct ways: (i) using a protocol recommended by MAKER2 authors for novel species and (ii) a protocol similar to the BRAKER2 training procedure (see 1.9). The predictions were generated on both repeat-masked and unmasked sequence.

Training by BRAKER2-like protocol (training on genes predicted by GeneMark-ES and at least partially supported by protein alignments) produced better prediction accuracy than training directly from protein alignments, which was the recommended MAKER2 protocol (Table S9).

MAKER2 gene predictions with GeneMark-ES and AUGUSTUS were more accurate than MAKER2 predictions which used the two gene finders along with SNAP, especially in terms of Sp values (Table S9). MAKER2 predictions with AUGUSTUS only were less accurate.

With the exception of *C. elegans*, the predictions on unmasked sequences (Table S10) showed an increase in prediction sensitivity and a decrease in specificity compared to the predictions on repeat-masked genome (Table S9). For *C. elegans*, we observed a decrease in both Sn and Sp when predictions were made on a masked genome. We attributed the decrease of the Sp value to MAKER2's hard-masking of all interspersed repeats (see section 1.9.1) which resulted in many predictions being corrupted due to repeat masking (11.9% of all annotated coding exons overlapped with sequences hard masked by MAKER2).

#### 2.2 AUGUSTUS training on the set of genes from the BUSCO families (the BUSCO genes)

BUSCO [12,13] pipeline has been frequently used for assessment of genome assembly completeness. It has been done by identifying a percentage of present in assembly single copy genes orthologous to genes in a clade specific BUSCO families. For the orthologues detection BUSCO has used AUGUSTUS-PPX [2] which has predicted genes in agreement with protein family profiles, in this case BUSCO protein families. The profiles of conserved OrthoDB protein families were provided by the authors of BUSCO. The BUSCO protocol has used one of the existing AUGUSTUS parameter sets as a starting point to predict genes that encode BUSCO proteins. Next, it used the identified genes to train AUGUSTUS for the target species. Subsequently, it has re-run the BUSCO protein detection, thus the BUSCO protocol has included the estimation of AUGUSTUS parameters for novel species.

We selected starting parameter sets that reflected the level of the taxonomic order exclusion: the *tomato* parameters for *A. thaliana*, the *ant* parameters for *D. melanogaster*, and the *schistosoma* parameters for *C. elegans.* It was not possible to exclude a target species from the precompiled BUSCO protein family profiles. In our experiments with the three model species, proteins of the target species and of close relatives therefore contributed to training AUGUSTUS on the target species in BUSCO.

We showed the *ab initio* gene prediction accuracy of AUGUSTUS trained by BRAKER2 for the three types of protein sets, as well as the *ab initio* gene prediction accuracy of AUGUSTUS trained on the 'BUSCO genes' (Table S11). In *A. thaliana*, the gene level F1 value achieved with BRAKER2 in the species exclusion scenario exceeded the accuracy achieved with BUSCO genes by ~7 percentage points. When all proteins of the same target order were excluded in BRAKER2, the accuracy still exceeded BUSCO by >1 percentage point. For *C. elegans*, the difference was larger. BRAKER2 exceeded BUSCO by 23 percentage points with the target species exclusion, by ~18 points and by ~20 points, for the family and order exclusion, respectively. For *D. melanogaster*, the BRAKER2 protocol achieved marginally higher accuracy than the BUSCO protocol (from 0.3 to 0.9 percentage points).
